# The Diffusion Exchange Ratio (DEXR): A minimal sampling of diffusion exchange spectroscopy to probe exchange, restriction, and time-dependence

**DOI:** 10.1101/2024.08.05.606620

**Authors:** Teddy X. Cai, Nathan H. Williamson, Rea Ravin, Peter J. Basser

## Abstract

Water exchange is increasingly recognized as an important biological process that can affect the study of biological tissue using diffusion MR. Methods to measure exchange, however, remain immature as opposed to those used to characterize restriction, with no consensus on the optimal pulse sequence(s) or signal model(s). In general, the trend has been towards data-intensive fitting of highly parameterized models. We take the opposite approach and show that a judicious sub-sample of diffusion exchange spectroscopy (DEXSY) data can be used to robustly quantify exchange, as well as restriction, in a data-efficient manner. This sampling produces a ratio of two points per mixing time: (i) one point with equal diffusion weighting in both encoding periods, which gives maximal exchange contrast, and (ii) one point with the same *total* diffusion weighting in just the first encoding period, for normalization. We call this quotient the Diffusion EXchange Ratio (DEXR). Furthermore, we show that it can be used to probe time-dependent diffusion by estimating the velocity autocorrelation function (VACF) over intermediate to long times (∼ 2 − 500 ms). We provide a comprehensive theoretical framework for the design of DEXR experiments in the case of static or constant gradients. Data from Monte Carlo simulations and experiments acquired in fixed and viable *ex vivo* neonatal mouse spinal cord using a permanent magnet system are presented to test and validate this approach. In viable spinal cord, we report the following apparent parameters from just 6 data points: *τ*_*k*_ = 17 ± 4 ms, *f*_*NG*_ = 0.71 ± 0.01, *R*_eff_ = 1.10 ± 0.01 *μ*m, and *κ*_eff_ = 0.21 ± 0.06 *μ*m/ms, which correspond to the exchange time, restricted or non-Gaussian signal fraction, an effective spherical radius, and permeability, respectively. For the VACF, we report a long-time, power-law scaling with ≈ *t*^− 2.4^, which is approximately consistent with disordered domains in 3-D. Overall, the DEXR method is shown to be highly efficient, capable of providing valuable quantitative diffusion metrics using minimal MR data.

---

Since its inception, the field of diffusion microstructural MR has developed many signal models to describe how features such as restriction (i.e., occupancy, size, and shape) [1–7] and processes such as exchange (i.e., permeability) [8–13] give rise to the diffusion MR signal. Despite advances in signal modelling, the development of experimental methods that can reliably disentangle these features have lagged behind. Largely, the field continues to rely on the classic pulsed-field gradient, spin echo (PGSE) method proposed by Stejskal & Tanner [14] in 1965. With PGSE, a.k.a. single diffusion encoding (SDE) wherein only the gradient amplitude is varied, restriction and exchange can manifest similarly. That is, variations in the signal behavior can be explained equally well by increased restriction or reduced exchange, or *vice versa*. The estimation of these parameters from SDE data is therefore degenerate, as discussed in refs. [15–21]. Due to this degeneracy, biological tissues that contain highly permeable compartment(s), such as gray matter (GM) [22], are difficult to characterize using SDE. The effects of restriction and exchange may coincide, meaning that estimated exchange times in GM — with some reports as fast as ≲ 10 ms [23–25] — are similar to a typical encoding period (≳ 10 ms). The robust quantification of exchange in such tissues requires the development of diffusion MR methods that go beyond SDE.

One approach is to extend the SDE framework by varying additional experimental parameters such as the diffusion time. This multi-dimensional data can then be fit to a signal model that includes both restriction and exchange [26–28], such as the models for neurite exchange imaging (NEXI) [29], or soma and neurite density imaging with exchange (SANDIX) [25, 30]. This SDE-based approach, while feasible, requires making *a priori* assumptions about the number of compartments and which compartments are exchanging. Exchange time estimates can vary substantially depending on the assumptions made (e.g., whether to include a “dot compartment” representing small, impermeable neurites [29] or whether to correct for the Rician noise floor [31]). As an example of this variability, neurite exchange time estimates in the human brain can vary by more than a factor of 2 (≈ 24 − 60 ms) depending on the model assumptions [31]. Furthermore, these methods do not fully address the issue of parameter degeneracy, as fit stability may remain difficult to achieve due to the large number of model parameters and small signal variations [25, 29, 31].

A potentially more robust approach to measure exchange is double pulsed-field gradient or diffusion encoding (DDE), which helps to resolve degeneracy by introducing additional experimental dimension(s) [20, 32]. In particular, diffusion exchange spectroscopy (DEXSY) is a DDE method proposed by Callaghan and Furó to separate water pools by their mobility and quantify the exchange between them [33]. The DEXSY experiment consists of two diffusion encoding periods along the same direction separated by a longitudinal storage period or mixing time, *t*_*m*_. Unlike the SDE-based approaches, DEXSY makes no assumptions about the number of compartments and their connectivity, but does assume Gaussian diffusion in all compartments by virtue of the Gaussian diffusion kernel. DEXSY has been found to produce accurate exchange parameters *in vitro* [34] and in phantom systems [35–37]. However, DEXSY is prohibitively data intensive in its original formulation and requires a well-sampled, two-dimensional grid of diffusion weightings per *t*_*m*_, making *in vivo* measurements infeasible. Clearly, experimental design optimization and data reduction are required to apply DEXSY to living systems. Progress has been made using classical techniques such as compressed sensing [38, 39] or constraints on the inversion [40], but few truly rapid methods that obviate the costly 2-D inversion have been proposed. These include filter exchange spectroscopy (FEXSY), proposed by Åslund *et al*. [41], and our own method [42, 43], which we expand upon here.

Restriction and exchange can also be viewed as giving rise to time-dependent diffusion [8, 44]. In parallel to the development of DEXSY and SDE-based signal models, another branch of diffusion MR theory and methodology emerged to measure the time-dependence of diffusion directly. Originating from the works of Stepišnik [45–47], these methods view the diffusion MR experiment in the frequency domain, wherein the spectrum of the (effective) gradient waveform *G*_eff_ (*t*) produces the diffusion weighting. Sequences with a sharp spectrum, such as a sinusoidal gradient oscillation in the time domain, can thus be swept to trace out the frequency-dependence of diffusion. This approach, called temporal diffusion spectroscopy (TDS) [48–50], has yielded promising results (e.g., refs. [50, 51]) but is limited in practice. High frequencies necessitate high slew rates and amplitudes whereas low frequencies can result in long echo times and *T*_2_ relaxation. TDS is confined to a somewhat narrow band of frequencies depending on the available gradient hardware and sample *T*_2_ (see Reynaud *et al*. [52] for review). And yet it is these difficult to access short- and long-time regimes that are best understood theoretically. In the short-time regime, universal scaling with 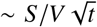 was found by Mitra *et al*. [53], where *S*/*V* is the surface-to-volume ratio (SVR). In the long-time regime, characteristic power law behaviors ∼ *t*^− *ϑ*^ were predicted by Novikov *et al*. [54] for what were termed “structural universality classes.”

These limitations arise in part due to the underlying theory of TDS. A frequency-domain representation implies the need for coherent oscillation, but in actuality any sequence will have some time- or frequency-domain weighting. Ning *et al*. [55] derived formulations of Stepišnik’s theory that remain in the time domain, and can thus be applied to general gradient sequences. For example, a given sequence can be viewed as a weighting of the ensemble mean-squared-displacement ⟨*r*^2^ (*t*)⟩ (MSD) by the autocorrelation function of *G*_eff_ (*t*). These signal representations enable the fitting of time-dependent diffusion using sequences that do not coherently oscillate, as shown by Cai *et al*. [56], for example. Viewing the DEXSY experiment through the lens of these representations may yield insights about how exchange and time-dependent diffusion are related.

In this work, we show that a particular sparse sub-sampling of DEXSY data can robustly measure exchange *and* restriction, as well as provide information about time-dependent diffusion from relatively little MR data. The sub-sampling consists of two DEXSY points that are equally diffusion-weighted, but one is maximally exchange-weighted while the other has little to no exchange-weighting. Our measurement approach is to take the quotient of these two points over various *t*_*m*_, and thus we call it the diffusion exchange ratio (DEXR) method. Compared to conventional SDE or TDS approaches, the DEXR method (i) overcomes degeneracy by isolating the estimation of exchange from the estimation of restriction, (ii) reduces the overall data requirements, and (iii) extends the range of sensitivity in the time domain by using the longitudinal mixing time *t*_*m*_ to shift the weighting, rather than the diffusion time in the transverse plane.

While a rapid measurement of exchange was described previously [42, 43], we aim here to provide a self-contained framework for DEXR experiments, and reiterate these previous findings. The novel contributions of this work are the means to estimate restriction parameters from the same data, the link to time-dependent diffusion, and validation of these concepts using Monte Carlo simulations. To limit the scope, we consider the case of static or constant gradients (SG) wherein the gradient amplitude *g* can be treated as a constant. Diffusion encoding is achieved by the SG spin echo (SG-SE) method, as in the first NMR measurement of diffusion [57]. In contrast to the way PG experiment are typically performed, SG experiments involve varying the time that spins spend in the effective gradient rather than the gradient amplitude. The application to PG, suitable for preclinical and clinical migration, is discussed briefly at the end of the manuscript.

The manuscript is organized as follows. After a description of the methods, we first discuss the signal behavior due to diffusion in the SG-SE experiment and propose a parsimonious signal model that incorporates both Gaussian and non-Gaussian signal behavior. Extending this model to SG-DEXSY, we show that just two points per *t*_*m*_ are sufficient to measure the (apparent) rate constant of exchange, *k*, as well as restriction (i.e., size and occupancy) with some reasonable assumptions. This forms the DEXR method. Optimal parameter selection is discussed. We then adopt an alternative view of this method in terms of time-dependent diffusion. We find that modifying *t*_*m*_ shifts a nearly point-wise sampling in the velocity autocorrelation function (VACF),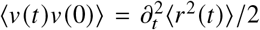. The DEXR method can thus be used to measure the VACF over a wide range of timescales (*t* ≈ 2 − 500 ms) compared to TDS with oscillating gradients. We support and validate our observations throughout with data acquired in fixed and viable *ex vivo* neonatal mouse spinal cords using a low-field, high-gradient system (*g* = 15.3 T/m), as well as data from Monte Carlo simulations in loosely packed, monodisperse spheres.

## 1. Materials and methods

### 1.1. Biological sample preparation

Spinal cords were extracted from Swiss Webster wild type mice (Taconic Biosciences, Rensselaer, NY, USA) via a ventral laminectomy under an animal protocol approved by the *Eunice Kennedy Shriver* National Institute of Child Health and Human Development Animal Care and Use Committee (Animal Study Proposal (ASP) # 21-025). Extracted spinal cords were bathed in low-calcium, high-magnesium artificial cerebrospinal fluid (aCSF, concentrations in mM: 128.35 NaCl, 4 KCl, 0.5 CaCl_2_ · 2H_2_O, 6 MgSO_4_ · 7H_2_O, 0.58 NaH_2_PO_4_ · H_2_O, 21 NaHCO_3_, 30 D-glucose).

Spinal cords were isolated together with the ventral roots and ganglia. In terms of size, the cords were roughly 15 × 1 × 1.5 mm (anterior–posterior length × lateral width × ventral–dorsal height). Data from fixed spinal cords and a single viable, *ex vivo* spinal cord are presented. For the fixed samples, fixation was performed immediately after dissection in 4% paraformaldehyde and the sample was left overnight at 4 °C. Fixative was then replaced with normal aCSF (same as before, but with 1.5 mM CaCl_2_ · 2H_2_O, 1 mM MgSO_4_ ·7H_2_O) 3 times over 2 days to remove residual paraformaldehyde before NMR measurements. For the viable sample, NMR measurements were performed immediately after dissection and the sample was kept alive in a wet/dry chamber with circulating aCSF bubbled with 95% O_2_, 5% CO_2_. All data is from spinal cords extracted between 1 − 4 days postnatal. Experiments were performed at a controlled temperature of 25 ± 0.2 °C, measured using a PicoM fiber optic temperature sensor (Opsens Solutions Inc, Québec, Canada) and controlled using an external water bath. Note that at this early stage of development, spinal cords predominantly consist of GM [58, 59], such that fast exchange is expected.

### 1.2. NMR hardware and methods

NMR experiments were performed on a low-field, single-sided, permanent magnet system: the PM-10 NMR-MOUSE (Magritek, Aachen, Germany) [60, 61]. This is an iron yoke magnet with a field strength that decays rapidly and roughly linearly with distance from the magnet’s surface. The active region is chosen as *B*_0_ = 0.3239 T, where the field is relatively uniform in a slice parallel to the magnet’s surface. The gradient arises from the linear decay of the static field, resulting in a strong SG of *g* ≈ 15.3 T/m, or *G* = *γg* ≈ 650 kHz/mm for the proton gyromagnetic ratio, *γ* ≈ 2.675 × 10^8^ s^− 1^ T^− 1^. The positioning of the magnet was controlled using a stepper motor with step size of 50 *μ*m. A custom-built solenoid was used as a transmit-receive radiofrequency (RF) coil. The coil was designed to fit the spinal cord(s) snugly with a high filling factor. Compared to flat coil designs, this gives a significant increase in the signal-to-noise ratio (SNR) [62, 23]. See refs. [23, 63] for a further description of the system and chamber. Diffusion measurements were performed using the standard SG-SE sequence with echo time 2*τ*, where *τ* is the spacing between the 90° and 180° RF pulses [64]. The 4-step phase cycle given in Table 2.2 of ref. [65] was used. For exchange measurements, an SG-DEXSY sequence was developed in Prospa (V3.22) that stores the signal at the time of echo formation. The 8-step phase cycle given in Appendix 5 of ref. [23] was used. When combined with unequal *b*-values for spoiling (*τ*_1_ ≠ *τ*_2_) this comprehensively suppresses off-resonance effects. The condition *τ*_1_ ≠ *τ*_2_ was achieved practically by offsetting *τ*_2_ by 0.013 ms in all SG-DEXSY measurements, avoiding exact parity. In Fig. 1, we show a diagram of the SG-DEXSY sequence and its modulation of the effective gradient *G*_eff_ (*t*) ∈ {0, − *γg*, +*γg*} by RF pulses.

**Figure 1:**
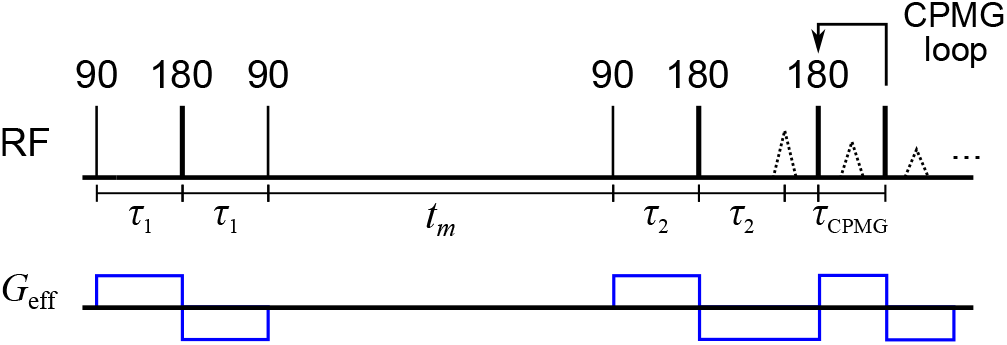
SG-DEXSY pulse sequence with timing parameters *τ*_1_, *τ*_2_, and *t*_*m*_. Signal is acquired in a CPMG loop with *τ*_CPMG_. The effective gradient *G*_eff_ (*t*) ∈ {0, − *γg*, +*γg*} and its modulation by RF pulses is shown below.

All experiments used hard RF pulses with pulse powers of − 22/16 dB (for 90°/180°-pulses) and duration ≈ 2 *μ*s. Pulses were driven by a 100 W amplifier (Tomco, Adelaide, Australia). For *g* = 15.3 T/m, this results in a sagittal slice of thickness Δ*z* ≈ 400 *μ*m. Measurements were performed with a Kea2 spectrometer (Magritek, Wellington, New Zealand). Phase correction was optimized at the start of the experiment such that signal in the real channel was maximized and signal in the imaginary channel was zero-mean. Data were acquired as signal from the real channel, summing over the echoes in a Carr-Purcell-Meiboom-Gill (CPMG) [66, 67] echo train with 2000 echoes and *τ*_CPMG_ = 12.5 *μ*s (see Fig. 1). This CPMG echo train acquisition is a common method to boost SNR in low-field experiments performed in an inhomogeneous **B**_0_ field [61, 65]. The repetition time (TR) was 2 s. Note that these NMR data were previously presented across refs. [23, 63], but are reanalyzed here to yield novel insights.

### 1.3. Monte Carlo simulations

Monte Carlo simulations were implemented in Julia 1.9.4. Monodisperse spheres with radius *R* = 0.95 *μ*m were placed in a 5 × 5 × 5 grid, equally-spaced, with centers 2 *μ*m apart and a minimum inter-sphere distance of 0.1 *μ*m. The spheres were situated inside of a 11 × 11 × 11 *μ*m box such that there is an empty surrounding space of 0.5 *μ*m in all directions. The overall intra-sphere volume fraction is calculated as ≈ 0.34. Simulations were performed with a time step of Δ*t* =2.5 × 10 ^− 4^ ms, and each walker step was a random sampling of the unit sphere 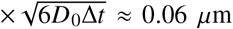, where *D*_0_ = 2.15 *μ*m^2^/ms was set to be near the measured diffusivity of water in aCSF at 25 °C [63]. To initialize the simulation, 10^4^ walkers were placed randomly and uniformly within the box. These simulation parameters are expected to yield low to moderate variability (≲ 5%) between repetitions with different random seeds [68]. Permeability and exchange were modelled using a small cross-over probability of 4 × 10^− 5^ upon collision with a sphere wall (and reflection otherwise). Specifically, walkers take a full step upon cross-over and otherwise experience a perfect (elastic) collision with the wall.

Gradients and phase accrual were simulated by having the isocenter through the central plane of the box and a gradient *g* = 15.3 T/m in the *x*-direction, consistent with the PM-10. The phase of each walker *ϕ*(*t*) was updated per step by *ϕ* (*t* + Δ*t*) = *ϕ* (*t*) + Δ*t*Δ*ω*, where *ϕ* 0 = 0, Δ*ω* is the local frequency offset given by Δ*ω* = (*x* − *x*_0_) *g*, with *x*_0_ = 5.5 *μ*m. The effect of 180° RF pulses was simulated as an instantaneous change in the sign of Δ*ω*. During the mixing time *t*_*m*_, Δ*ω* was set to 0. A reflecting, rather than periodic boundary condition was used at the edge of the box to avoid issues related to changes in Δ*ω* upon exiting the domain. Finally, the signal was calculated by taking the real part of ⟨exp (*iϕ*)⟩, where ⟨·⟩ represents ensemble averaging. Relaxation effects were not included but could be implemented as the subject of future work.

## 2. Diffusion in the SG-SE experiment

Before addressing exchange and SG-DEXSY, we must first consider the SG-SE experiment with echo time 2*τ* and gradient amplitude *g*. The SG-SE is the basic experimental paradigm used here, as the implemented SG-DEXSY sequence consists of two SG-SE blocks (see Fig. 1). According to Hürlimann *et al*. [69] (c.f., refs. [70–73] for review), one can roughly separate the SG-SE signal behavior due to diffusion into three regimes: Gaussian, and two non-Gaussian regimes. In the Gaussian regime, spin isochromats or simply “spins” are not significantly impeded by barriers and diffusion is effectively free, resulting in Torrey’s [74] well-known expression for the normalized signal attenuation: *S*/*S*_0_ = exp (− *bD*_0_), where *b* = (2/3) *γ*^2^*g*^2^*τ*^3^, *γ* is the proton gyromagnetic ratio, and *D*_0_ is the self-diffusion coefficient of water.

### 2.1. Asymptotic regimes

In the non-Gaussian regime(s), spins are impeded and the limiting signal behavior takes on a much slower decay (n.b., the term non-Gaussian is used here to refer to the distribution of spin displacements and restriction by barriers in general, rather than to the phase distribution). The form of the signal decay depends on the relationship between characteristic length scales: (i) the diffusion length 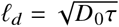, which is the typical distance travelled by spins during each gradient application, (ii) the structural length *𝓁*_*s*_, which defines the confinement dimension along the gradient axis (e.g., pore diameter), and (iii) the dephasing length *𝓁*_*g*_ = (*D*_0_/*γg*) ^1/3^, which is the distance that two spins must travel to de-correlate their phase by *π* radians. Any one of these length scales being much shorter than the others gives rise to a different asymptotic regime of signal behavior. The Gaussian regime arises when *𝓁*_*d*_ is much shorter than *𝓁*_*s*_. In terms of these length scales, we have that:

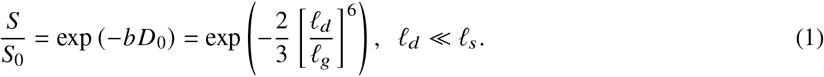

The “motional averaging” or “motional narrowing” regime arises when *𝓁*_*s*_ is the shortest of the three length scales such that spins experience only a limited range of frequencies over time [3, 75]. Exact solutions were given by Neuman [5] in the case of simple, impermeable domains. For spheres of radius *R*:

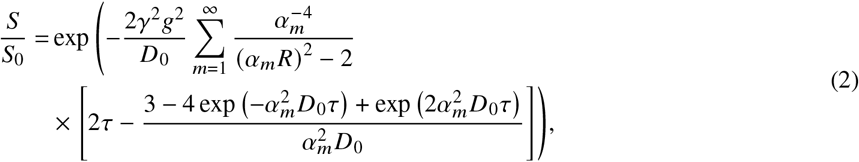

where *α*_*m*_ is *m*^th^ root of

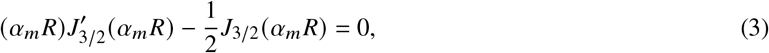

and *J* represents a Bessel function of the first kind. The first 5 roots are sufficient to obtain a good approximation for short diffusion times (≲ 1 ms) and small radii (≲ 1 *μ*m), and are given by *α*_*m*_*R* = [2.0815, 5.940, 9.206, 12.405, 15.579]. In the limit of large *𝓁*_*d*_, the above expression simplifies to:

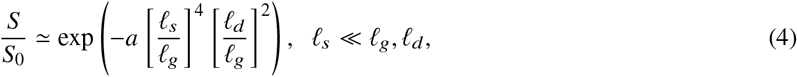

where *a* is a geometry-dependent prefactor (e.g., for spheres, *a* = 1/175 and *𝓁*_*s*_ = 2*R*). The “localization” regime arises when *𝓁*_*g*_ is shortest. In this regime, signal localized near barriers within a distance of *𝓁*_*g*_ persists whereas signal deeper within the structure dephases due to being able to displace a distance *𝓁*_*d*_ > *𝓁*_*g*_ within *𝓁*_*s*_. The signal behavior in this regime was first described by Stoller *et al*. [76]. To a first-order approximation, the asymptotic signal behavior at large *𝓁*_*d*_ (ignoring permeability) is given by [72, 76–78]

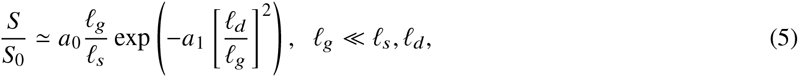

where *a*_0_ is a geometry-dependent prefactor (e.g., *a*_0_ = 5.8841 for parallel plates), and *a*_1_ = 1.0188 is a universal prefactor. While the signal behavior in these non-Gaussian regimes is complicated and exact expressions are either unwieldy or not available, the different scaling behaviors in terms of *𝓁*_*d*_, *𝓁*_*g*_, and *𝓁*_*s*_ are clear.

### 2.2. A parsimonious ensemble signal model

As noted by Grebenkov [72, 79], these scaling behaviors yield a simple, dichotomous view of the Gaussian and non-Gaussian regimes for the SG-SE experiment. Consider that the *b*-value is proportional to *τ*^3^ or 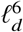. Since *g* is fixed in this case, *𝓁*_*g*_ is constant and *τ* is the only parameter being varied. The Gaussian and non-Gaussian regimes are contrasted by their (*𝓁*_*d*_/*𝓁*_*g*_)^6^ vs. (*𝓁*_*d*_/*𝓁*_*g*_)^2^ scaling, respectively — see Eqs. (4) and (5). This ratio can be seen as the controlling feature of the SG-SE experiment, so we define the following dimensionless parameter, *ρ*:

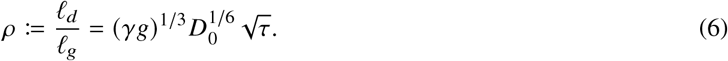

Equivalently stated, signal in the Gaussian regime decays with *b* ∝ *ρ*^6^ whereas the non-Gaussian signal decays much more slowly with *b*^1/3^ ∝ *ρ*^2^ [79, 80]. We illustrate this dichotomy in Fig. 2, plotting the normalized signal decay from Eq. (2) for spheres of radii *R* = 0.4 − 1 *μ*m in comparison to free diffusion as a function of *ρ*^2^. As *𝓁*_*d*_ increases and *ρ*^2^ ≫ 1, the signal behavior for spheres quickly approaches the asymptotic, linear behavior predicted by Eq. (4). The parameter values are chosen to correspond to the PM-10 system at room temperature: *D*_0_ = 2.15 *μ*m^2^/ms, *g* = 15.3 T/m. These values are used throughout, though we stress that by expressing the signal w.r.t. powers of *ρ*, the observations can be generalized to other SG systems with different attributes. Also plotted is the localized signal decay from Eq. (5), which may become relevant as *ρ*^2^ increases, at least for larger *𝓁*_*s*_ = 2*R* ≫ *𝓁*_*g*_.

**Figure 2:**
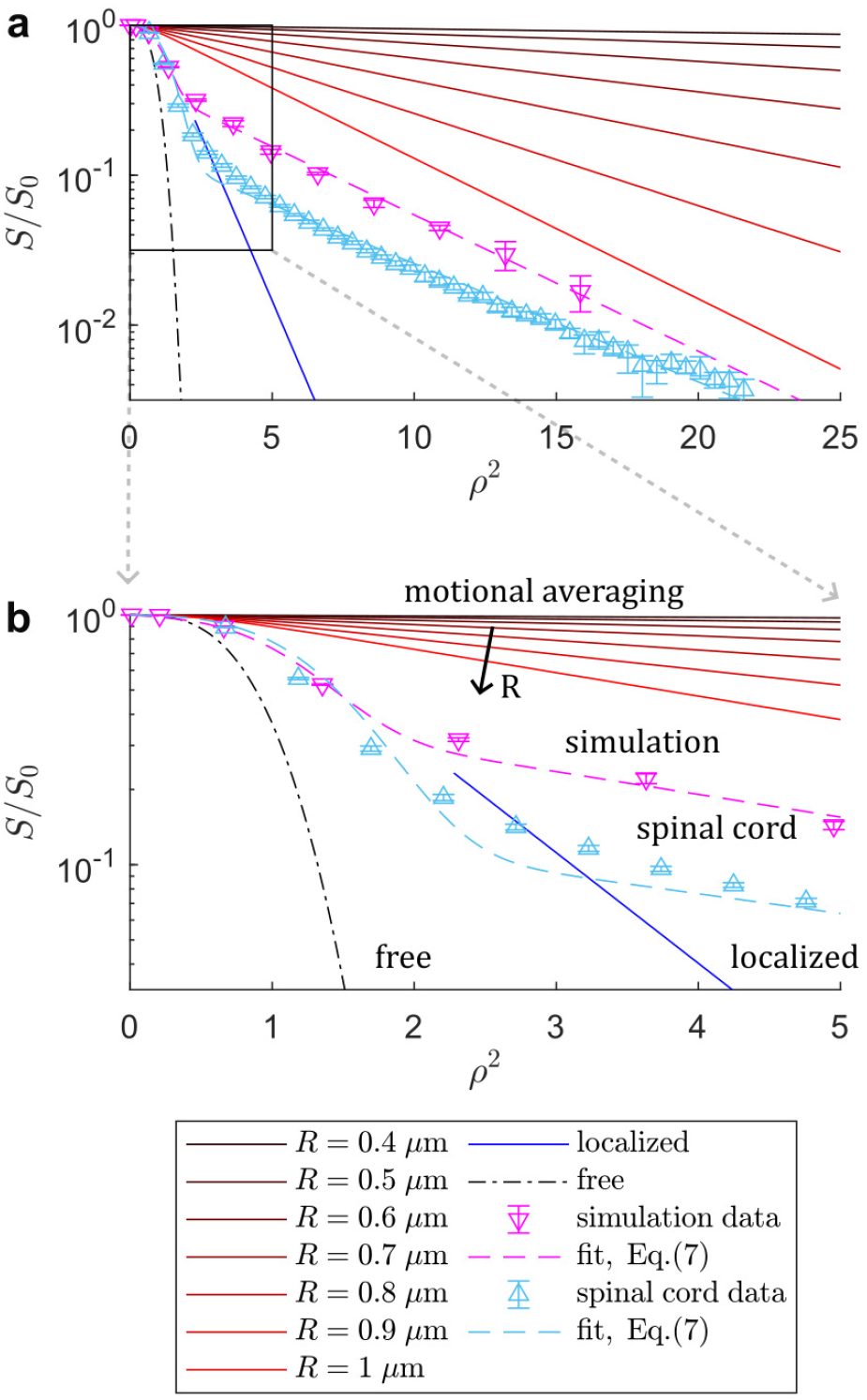
Comparison of the SG-SE signal behavior in different regimes and in simulation and fixed spinal cord data w.r.t. *ρ*^2^ = (*𝓁*_*d*_/*𝓁*_*g*_)^2^ ∝*b*^1/3^. **(a)** Signal curves *S*/*S*_0_ plotted on a log-axis for the free (dash-dot line), motionally-averaged (red to black lines), and localized (blue) regimes compared to data from Monte Carlo simulations (magenta) and data acquired in fixed spinal cord (cyan). Curves are plotted up to *ρ*^2^ = 25 using *D*_0_ = 2.15 *μ*m^2^/ms, *g* = 15.3 T/m, which gives a dephasing length *𝓁*_*g*_ ≈ 806 nm. Motionally-averaged signal is plotted for spherical radii from *R* = 0.4 − 1 *μ*m or from *𝓁*_*s*_ = 2*R* ≈ 1 − 2.5 *𝓁*_*g*_, summing up to the first 5 roots in Eq. (2). Note the rapid approach of Eq. (2) towards the asymptotic, linear behavior predicted by Eq. (4) as *ρ*^2^ exceeds ≈ 2, or *ρ* ≳ 1.4. Localized signal is plotted only for *𝓁*_*d*_ > 1.5 *𝓁*_*g*_ using the prefactor *a*_0_ = 5.8441, see Eq. (5). For the Monte Carlo simulation data, error bars indicate ± 1 SD from 3 repetitions with different random seeds. For the spinal cord data, error bars indicate 95% confidence intervals estimated by bootstrapping 43 repetitions on the same sample. Fits to Eq. (7) yield *f*_*I*_ ≈ 0.44, ⟨*c*_*E*_⟩ ≈ 0.40, ⟨*c*_*I*_⟩ ≈ 0.21 for the simulation data and *f*_*I*_ ≈ 0.16, ⟨*c*_*E*_⟩ ≈ 0.26, ⟨*c*_*I*_⟩ ≈ 0.18 for the spinal cord data. **(b)** Zoomed plot up to *ρ*^2^ = 5, highlighting the transition from Gaussian signal behavior to the characteristic non-Gaussian signal decay that is linear on this axis of *ρ*^2^ ∝ *b*^1/3^. Note the deviation from the fit in the spinal cord data, suggestive of potentially distributed non-Gaussian compartments.

The picture is more complicated in heterogeneous environments such as biological tissue, which may be hierarchically organized, and wherein there may be a range of *𝓁*_*s*_ values present. In such samples, all three of these regimes can arise within different microenvironments, and the ensemble signal resists characterization by any one of the signal expressions. Nonetheless, according to the dichotomous view above, the non-Gaussian signal can be lumped into some *effective* decay with *ρ*^2^, irrespective of the actual distribution of *𝓁*_*s*_ and the mixture of motionally-averaged and localized signal that may arise as a result. The ensemble signal can be approximated as a Gaussian signal fraction decaying with *ρ*^6^ and a non-Gaussian fraction decaying with *ρ*^2^, as suggested by Cai *et al*. [81], and which is similar in principle to the combined hindered and restricted (CHARMED) model [82]. Ignoring exchange for the time being, we can write the following quasi-biexponential model:

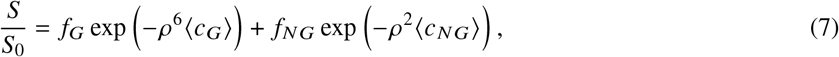

where *f*_*G*_ represents the Gaussian fraction (e.g., the occupancy fraction of the extracellular space, or ECS), *f*_*NG*_ represents the restricted, non-Gaussian fraction (e.g., the intracellular space, or ICS), *f*_*G*_+*f*_*NG*_ = 1, and ⟨*c*_*G*_⟩, ⟨*c*_*NG*_⟩ are dimensionless decay constants w.r.t. *ρ*^6^ and *ρ*^2^, respectively, where ⟨·⟩ represents signal-weighted ensemble averaging. For free diffusion, ⟨*c*_*G*_⟩= 2/3, with smaller values indicating hindered diffusion with an apparent diffusivity *D*_app_ given by

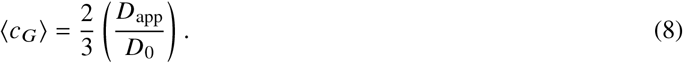

The non-Gaussian decay ⟨*c*_*NG*_⟩ can be viewed as arising from some effective structure size. In the case of motional averaging within spheres, we obtain from Eq. (4),

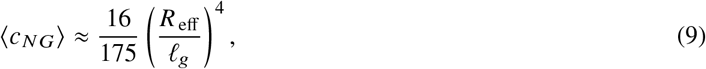

which is valid for large *ρ* ≫ 1 (*𝓁*_*d*_ ≫ *𝓁*_*g*_, *𝓁*_*s*_), and where *R*_eff_ is an effective spherical radius, that by volume weighting may take the form *R*_eff_ = (⟨*R*^7^⟩/⟨*R*^3^⟩)^1/4^ [83]. Note that in this model, the signal fractions *f*_*G*_, *f*_*NG*_ do not represent volume fractions *per se*. Rather, they represent the proportion of signal that appears to undergo Gaussian vs. non-Gaussian signal decay. Water within some large structure with *𝓁*_*s*_ ≫ *𝓁*_*g*_, for instance, may include both Gaussian and non-Gaussian decay, with some signal that dephases with *ρ*^6^ and some that becomes localized and dephases with *ρ*^2^ [79, 78].

In Fig. 2, we show that both simulated and real SG-SE data measured in fixed spinal cord fit well to Eq. (7). For the simulation data, a fit to the mean yields: *f*_*NG*_ ≈ 0.44, ⟨*c*_*G*_⟩ ≈ 0.40, ⟨*c*_*NG*_⟩ ≈ 0.21. For the spinal cord data: *f*_*NG*_ ≈ 0.16, ⟨*c*_*G*_⟩ ≈ 0.25, ⟨*c*_*NG*_⟩ ≈ 0.18. The ⟨*c*_*NG*_⟩ obtained from simulation data yields *R*_eff_ ≈ 1.0 *μ*m from Eq. (9), which roughly agrees with the actual radius *R* = 0.95 *μ*m, though the fitted *f*_*NG*_ ≈ 0.44 overestimates the intra-sphere volume fraction of ≈ 0.34. This may be because the space between spheres can appear to be restricted rather than hindered — consider that at its narrowest, the inter-sphere spacing is 0.1 *μ*m (see Methods). Another confounding effect is that as *ρ* and *𝓁*_*d*_ increase, exchange during the encoding will also increase, which may reduce the effect of restriction (i.e., increase ⟨*c*_*NG*_⟩ and/or reduce *f*_*NG*_). As discussed, the estimation of restriction and exchange parameters is degenerate with SDE. In the spinal cord data, there is notable deviation from the fit around the transition at *ρ*^2^ ≈ 2 − 4 (see Fig. 2b). This deviation could be explained by the different rates that the (potentially) numerous non-Gaussian signal pools approach the limiting behavior whence *ρ*^2^ scaling emerges. A related issue is that for *ρ* ≲ 1, the non-Gaussian signal is not yet well-described by a simple scaling with *ρ*^2^ and further terms that were truncated to arrive at Eq. (4) are needed to explain the signal [5]. Due to these issues, the fit parameters to Eq. (7) should be treated as apparent and non-quantitative.

Despite its limitations, Eq. (7) is seen to be a good empirical signal model for systems that contain both Gaussian and non-Gaussian signal populations, and can fit the data well across a wide range of *ρ*^2^ values. Importantly, the model captures the distinct scaling behaviors that differentiate the Gaussian and non-Gaussian signal pools, and provides a starting point for our modelling of the SG-DEXSY signal.

## 3. Exchange and restriction in the SG-DEXSY experiment

How does this signal model relate to exchange and the SG-DEXSY experiment with parameters *τ*_1_, *τ*_2_, *t*_*m*_? Note that by exchange, we refer specifically to barrier-limited exchange, when molecules typically diffuse across a structure many times before exiting. This can be more formally stated using a permeability length: *𝓁*_*k*_ = *D*_0_/*k* (see Novikov [13], c.f., Grebenkov [84]), where *κ* is the permeability with units of length per time. The permeability length can be seen as an effective membrane thickness or as a competition between diffusive and barrier-limited kinetics. If exchange is limited by the time to diffuse to the barrier (*𝓁*_*k*_≪*𝓁*_*s*_) the effect of exchange is indistinguishable from hindered diffusion. The long-time limit is rapidly reached and Gaussian diffusion is recovered. Barrier-limited exchange is observable only when *𝓁*_*k*_ ≫ *𝓁*_*s*_ and *𝓁*_*d*_ ≫ *𝓁*_*s*_. This means that motionally-averaged or localized signal *must* be present; the barrier-limiting condition is tantamount to non-Gaussian signal behavior. In this case, exchange can be modelled with a first-order rate constant, *k* [9, 10, 84]:

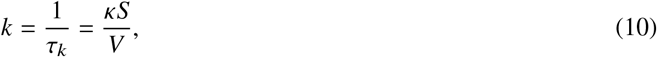

where *τ*_*k*_ is the corresponding exchange time (i.e., a mean pore residence time) and *S*/*V* is the SVR. As an aside, we point out that any signal model that characterizes the confined signal using a hindered diffusivity yet also fits a first-order exchange rate, such as the Kärger model [9, 10] or indeed the original DEXSY model with *P* (*D*_1_, *D*_2_)[33], does not correctly describe the SG experiment.

With the preceding remarks on diffusion in mind, the SG-DEXSY experiment can be conceptualized as follows: (i) spins undergo an initial diffusion encoding with echo time 2*τ*_1_ that separates microenvironments into Gaussian and non-Gaussian regimes by their degree of dephasing, (ii) the signal in these environments then mix during the longitudinal storage period *t*_*m*_, wherein exchange *out* of the non-Gaussian microenvironments is barrier-limited, and (iii) a second encoding with 2*τ*_2_ dephases the exchanging signal, resulting in exchange-weighted contrast in the measured echo intensity, *S*. For this experiment to work, several conditions must be met.

### 3.1. Sensitivity to exchange

Firstly, exchange during *t*_*m*_ must be detectable. That is, exchange must not proceed so quickly that a steady state is reached during the encoding itself: *τ*_*k*_ ≫ 2*τ*_1_. Another condition is that the signal must not fully decay by *T*_1_ relaxation, (i.e., *T*_1_ ≫ *t*_*m*_). There must also be significant contrast between the Gaussian and non-Gaussian signals. According to the above conception of the SG-DEXSY experiment, the sensitivity to exchange is proportional to the difference in signal decay between these regimes, with exchange having the greatest effect when they are maximally separated. Practically, this translates to parameters in the regime of *bD*_0_ ≳ 2 (i.e., *ρ* ≳ 1.2) such that the initial encoding greatly dephases the Gaussian signal while preserving the non-Gaussian signal. This is similar to the notion of an efficient “filtering” value of *b*_1_ in FEXSY [41, 85]. This condition also ensures that the non-Gaussian signal can in fact be described as scaling with *ρ*^2^ (see Fig. 2b).

It is important to note that the range of structure sizes *𝓁*_*s*_ for which exchange is being probed is determined by the chosen *𝓁*_*d*_ and *𝓁*_*g*_. Consider that the available *g* (and *𝓁*_*g*_) determines how large *τ* (and *𝓁*_*d*_) must be to achieve signal separation and *ρ*^2^ > 1. Subsequently, *𝓁*_*d*_ dictates what values of *𝓁*_*s*_ result in restriction or non-Gaussian signal (*𝓁*_*s*_ ≪ *𝓁*_*d*_). Thus, the measurement is sensitive to exchange *out* of structures for which *𝓁*_*s*_ ∼ *𝓁*_*g*_ ≈ 0.8 *μ*m ≲ *𝓁*_*d*_. Consider too the condition about exchange during the encoding (*τ*_*k*_ ≫ 2*τ*_1_). The choice of 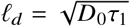 dictates what values of *τ*_*k*_ can be measured via 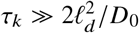. Thus, the sensitivity to *τ*_*k*_ and *𝓁*_*s*_ has a multi-faceted dependence on *𝓁*_*g*_. In general, higher *g* enables the measurement of faster exchange out of smaller structures [84]. This dependence may explain in part the large range of reported exchange times in the literature for tissues with heterogeneous microstructure, which can vary by more than an order of magnitude in ostensibly similar tissue [86].

### 3.2. Optimal parameter selection

What value of *ρ* exactly maximizes sensitivity to exchange? Put another way, what is the maximal signal difference Δ*S*/*S*_0_ between non-Gaussian and Gaussian signal w.r.t. *ρ*? As a first comparison, we look at Δ*S*/*S*_0_ for motional averaging vs. free diffusion, or Eq. (2) vs. Eq. (1), shown in Fig. 3. We plot only *ρ* > 1, keeping in mind that the signal model in Eq. (7) is only valid for *ρ* ≫ 1 whence the *ρ*^2^ scaling of the non-Gaussian regime(s) emerges. The maximum values cluster around *ρ* ≈ 1.3 − 1.5 for the chosen values of 2*R* ≲ *𝓁*_*g*_, with smaller radii leading to a larger optimum *ρ*. We also plot Δ*S*/*S*_0_ between the two decay terms exp (− *ρ*^6^ ⟨*c*_*G*_⟩) and exp(− *ρ*^2^ ⟨ *c*_*NG*_⟩) for the fits of Eq. (7) to simulation and spinal cord data, shown earlier in Fig. 2. For these fitted parameters, the maximal value of Δ*S*/*S*_0_ is smaller due to the hindered model of the Gaussian signal. This also results in a shift of the optimal *ρ* to the right. Nonetheless, the optima lie near the upper end of the values predicted for spheres, at *ρ* ≈ 1.45 and 1.55 for simulation and spinal cord data, respectively. In general, *ρ* ≳ 1.4 (or *ρ*^2^ ≳ 4, see again Fig. 2b) is a reasonable heuristic to achieve separation between the Gaussian and non-Gaussian signal without prior knowledge of *𝓁*_*s*_. This value of *ρ* corresponds to *τ* ≈ 0.59 ms and *b* ≈ 2.3 ms/*μ*m^2^ for *D*_0_ = 2.15 *μ*m^2^/ms and *g* = 15.3 T/m. Note that for *ρ* ≈ 1.3 − 1.5, localized signal is not expected because *𝓁*_*d*_ is only moderately larger than *𝓁*_*g*_, and the more straightforward interpretation of the non-Gaussian decay as arising from some effective spherical radius, ⟨*c*_*I*_⟩ = 16/175 (*R*_eff_/*𝓁*_*g*_)^4^ as in Eq. (9), is likely to be valid.

**Figure 3:**
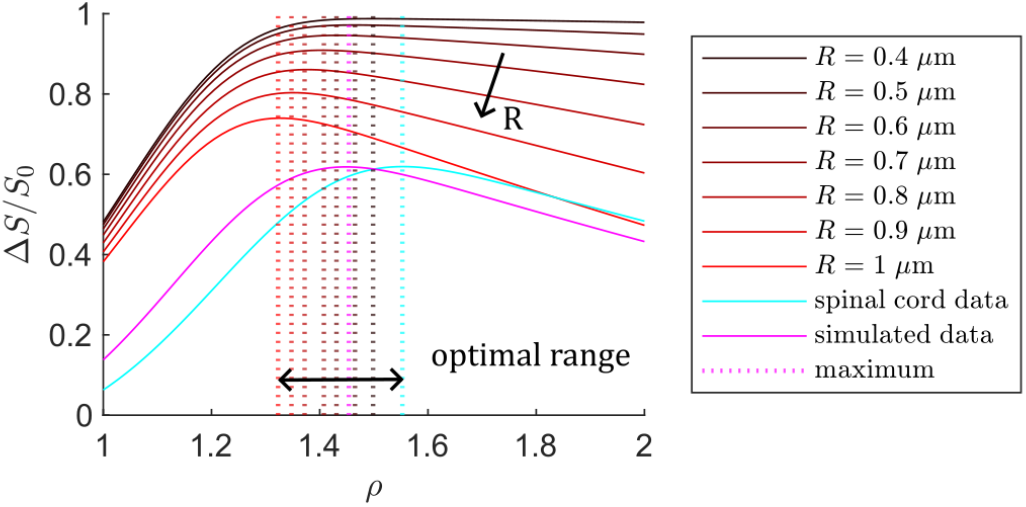
Signal difference Δ*S*/*S*_0_ between non-Gaussian and Gaussian signal decay plotted as a function of *ρ* = *𝓁*_*d*_/*𝓁*_*g*_ for motional averaging in spheres of radii *R* = 0.4 − 1 *μ*m using Eq. (2), compared to free diffusion, or Eq. (1). The difference between the components of the fits to Eq. (7) for simulated (magenta) and spinal cord (cyan) data shown in Fig. 2 are also plotted, i.e., Δ*S*/*S*_0_ is calculated as the difference between the terms exp(− *ρ*^2^ ⟨*c*_*NG*_ ⟩) and exp(− *ρ*^6^ ⟨*c*_*G*_ ⟩). For these data, Δ*S*/*S*_0_ is smaller and the maximum is farther to the right due to hindered diffusion. Overall, the optimal range to maximize Δ*S*/*S*_0_ is about *ρ* ≈ 1.35 − 1.55, with *ρ* ≳ 1.4 as a heuristic.

There are, however, *two* values of *ρ* in SG-DEXSY, with *ρ*_1_ and *ρ*_2_ corresponding to *τ*_1_ and *τ*_2_. The above answers what value of *ρ*_1_ is optimal, but what of *ρ*_2_? One might guess that holding *ρ*_1_ = *ρ*_2_ is optimal, again maximally separating the Gaussian and non-Gaussian regimes in the second encoding. This is in fact the case, as was shown in our previous work [42, 81] and by others [21, 87]. We reiterate this result by extending Eq. (7) to SG-DEXSY, writing the signal as arising from four signal fractions:

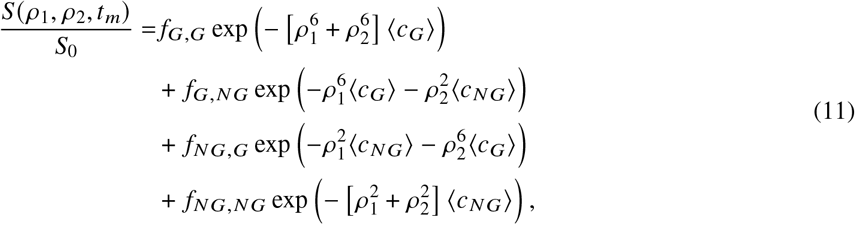

where *f*_*NG,G*_ represents the signal fraction that exchanges from a non-Gaussian to a Gaussian regime during *t*_*m*_ and so forth for *f*_*G,G*_, *f*_*G,NG*_, *f*_*NG,NG*_ (which sum to 1). Although the model appears to ignore exchange between the microenvironments that may comprise *f*_*G*_ and *f*_*NG*_ — i.e., it looks only at exchange between two bulk pools — we argue that if a (detailed) mass balance holds, then the ensemble-averaged decay constants ⟨*c*_*G*_⟩, ⟨*c*_*NG*_⟩ will not change with *t*_*m*_ and the pools can be treated as decaying identically in both encodings. Therefore, further components are not necessary to explain the signal behavior. Mass balance also implies that the exchanging signal fractions *f*_*G,NG*_, *f*_*NG,G*_ are equal such that we can define a total exchanging signal fraction

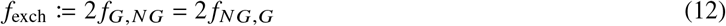

and rewrite the previous expression in terms of the equilibrium signal fractions, *f*_*G*_ and *f*_*NG*_, and *f*_exch_:

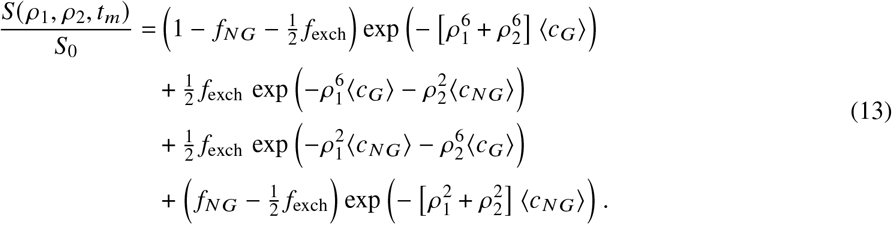

Note that the maximum possible value of *f*_exch_, which we will call *f*_exch, ss_ for steady-state, is given as a direct result of mass balance by

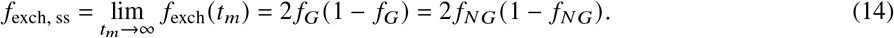

We also argue that exchange during the encoding period is implicitly accounted for in Eq. (13) because signal that exchanges partway through an encoding can nonetheless be modelled by some combination of the terms above. Incorporating a model of the intra-encoding exchange such as the Kärger model [9, 10] is not necessary, though such exchange may result in *f*_exch_ > 0 at *t*_*m*_ = 0.

In Figs. 4a and b, we plot signal contour maps generated by substituting the parameters obtained by fitting Eq. (7) to the SG-SE spinal cord data (see again Fig. 2) into Eq. (13). Contour maps are plotted for *ρ*_1_, *ρ*_2_ ≥ 1 and for several values of *f*_exch_ = [0.02, 0.13, 0.27], where the largest value *f*_exch_ = 2 *f*_*NG*_ (1 − *f*_*NG*_) ≈ 0.27 corresponds to near full signal turnover. In the rightmost plot, we look at the signal contrast Δ*S*/*S*_0_ due to exchange by taking the difference between the higher *f*_exch_ cases and the *f*_exch_ = 0.02 case. These difference maps indicate clearly that the maximal contrast is obtained when *ρ*_1_ = *ρ*_2_ and confirm the result shown in Fig. 3: that *ρ* ≈ 1.55 is optimal for obtaining exchange contrast with these parameters. Moving away from parity results in less contrast, indeed, none along the axes where *ρ*_1_ or *ρ*_2_ = 0. It is also clear that the contrast roughly doubles as *f*_exch_ doubles, indicating proportionality of this midpoint in the domain with *f*_exch_.

**Figure 4:**
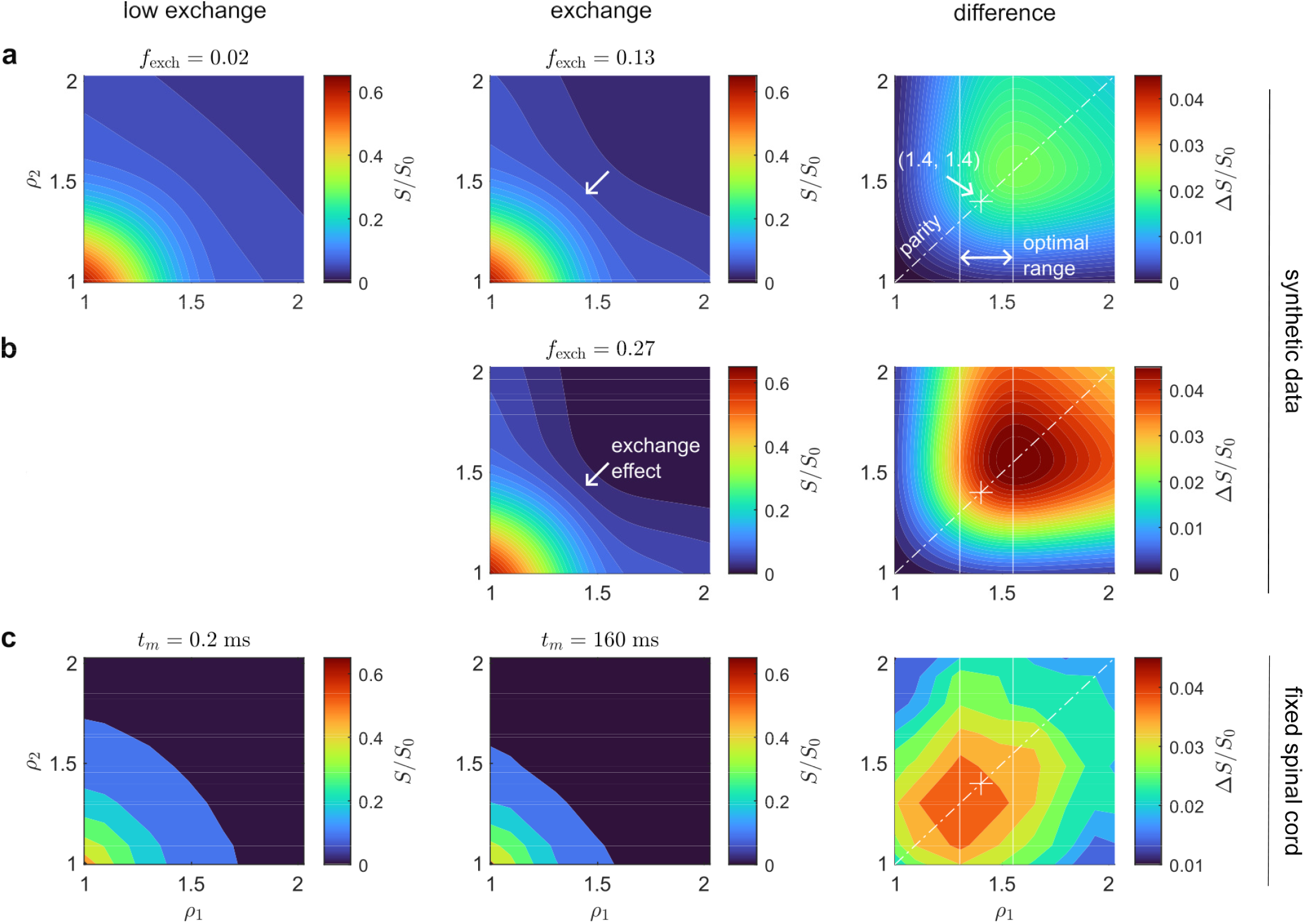
Signal contour and difference maps between low and high exchange cases. **(a)** Plots for synthetic data generated using Eq. (13) and the fit parameters obtained by fitting Eq. (7) to the SG-SE spinal cord data in Fig. 2: *f*_*NG*_ ≈ 0.16, ⟨*c*_*G*_⟩ ≈ 0.25, ⟨*c*_*NG*_⟩ ≈ 0.18. Exchanging signal fractions *f*_exch_ = [0.02, 0.13, 0.27] are compared, where *f*_exch, ss_ ≈ 0.27. Exchange is seen to produce an inwards curvature in the signal contours around *ρ*_1_, *ρ*_2_ ≳ 1.25 (see middle panel). The difference map Δ*S*/*S*_0_ indicates that *ρ*_1_ = *ρ*_2_ ≈ 1.55 produces the most exchange contrast, which agrees with the optimum and range identified in Fig. 3. The parity axis *ρ*_1_ = *ρ*_2_ is marked with a dash-dot line. The heuristic optimum of *ρ*_1_ = *ρ*_2_ = 1.4 is marked by a cross. **(b)** The same plots as part (a) but using the maximal *f*_exch_ = *f*_exch, ss_ ≈ 0.27. Note that the exchange contrast Δ*S*/*S*_0_ roughly doubles as *f*_exch_ doubles between (a) and (b), proportional with the increase in *f*_exch_. The peak value of Δ*S*/*S*_0_ increases from ≈ 0.023 to ≈ 0.046 (see color bar values). **(c)** The same plots for data acquired in fixed spinal cord in a 6 × 6 grid at *ρ*_1_ ≈ *ρ*_2_ = [1.09, 1.30, 1.49, 1.65, 1.80, 1.93] (recall that *τ*_2_ ≠ *τ*_1_, see Methods). In these data, the optimal point is shifted towards a smaller *ρ*_1_ = *ρ*_2_ ≈ 1.37 than in parts (a) or (b), and is also of a smaller peak amplitude than part (b), with a maximal Δ*S*/*S*_0_ ≈ 0.037. This may be due to the model being an incomplete description of the distributed non-Gaussian microenvironments in tissue (see again the fit deviations in Fig. 2b) and/or larger compartments with a smaller expected optimum but larger volume dominating the exchange contrast (see Fig. 3). Nonetheless, the heuristic *ρ*_1_ = *ρ*_2_ = 1.4 remains a good choice. Despite the coarse sampling of this data and the shift in optimum, the qualitative similarity in shape and character to part (b) is evident.

In Fig. 4c, we show analogous plots for data acquired in fixed spinal cord data over a 6 × 6 grid of *ρ*_1_ ≈ *ρ*_2_ = 1.09, 1.30, [1.49, 1.65, 1.80, 1.93] at a short *t*_*m*_ = 0.2 ms vs. a long *t*_*m*_ = 160 ms. At this long *t*_*m*_, exchange is expected to have reached the steady state, *f*_exch, ss_. Despite the coarse sampling of this data, the finding that *ρ*_1_ = *ρ*_2_ is optimal remains clear and the qualitative similarity to part (b) is evident. While the optimum is shifted slightly towards a smaller *ρ* ≈ 1.37, this may be due to the deviation from the fit around these values of *ρ*, which can be seen

Fig. 2b. Another explanation is that larger compartments dominate the exchange contrast due to their greater volume fraction (see the trend with *R* in Fig. 3). Regardless, *ρ*_1_ = *ρ*_2_ = 1.4 is shown to be a good heuristic and is marked by a cross in all of the difference maps.

### 3.3. The curvature method

Our goal though is not merely to obtain maximal exchange weighting, we also wish to isolate the effect of exchange such that the fitting of a highly parameterized model such as Eq. (13) is not necessary to estimate *τ*_*k*_. We should thus ask what set of SG-DEXSY points yields contrast due to exchange independent of other effects. We previously showed in Cai *et al*. [42] that by holding the sum of *b*-values — *b*_*s*_ = *b*_1_ + *b*_2_ — constant, we can isolate exchange from non-exchanging, Gaussian diffusion (an idea inspired by Song *et al*. [88]). We further showed that the curvature along an axis of constant *b*_*s*_ (i.e., along the difference axis *b*_*d*_ = *b*_1_ − *b*_2_) is proportional to *f*_exch_ if the exchanging microenvironment(s) can be adequately modelled with an apparent diffusivity, i.e., decaying as exp (− *bD*_app_). From Eq. (9) in Cai *et al*. [42], a minimal measurement of *f*_exch_ at a given *t*_*m*_, assuming two sites with diffusivities *D*_*E*_ > *D*_*I*_, is

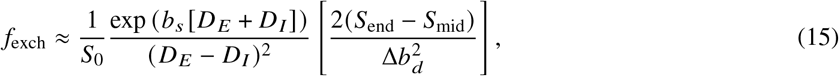

where Δ*b*_*d*_ is a step-size in *b*_*d*_ as close to *b*_*s*_ as possible, *S*_end_ corresponds to the signal when (*b*_1_, *b*_2_) = (Δ*b*_*d*_, *b*_*s*_ − Δ*b*_*d*_), *S*_mid_ corresponds to 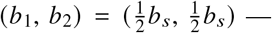., the point along the parity axis with maximal exchange weighting — and the bracketed term on the right-hand-side is a finite difference approximation of the curvature in *S* w.r.t. *b*_*d*_ about *b*_*d*_ = 0, taking advantage of the symmetry across the parity axis. The general approach is visually supported by the rightmost column of Fig. 4, which shows that (*S*_end_ − *S*_mid_)/*S*_0_ takes the difference between a point with almost no exchange weighting along *ρ*_1_ or *ρ*_2_ ≈ 0 (i.e., Δ*b*_*d*_ ≈ *b*_*s*_) and a point with maximal weighting along parity *ρ*_1_ = *ρ*_2_, thereby isolating exchange. The two points are notated as such because *S*_mid_ corresponds to a midpoint in the domain and *S*_end_ corresponds to an endpoint along the marginal axis.

As discussed in the previous section, however, a model such as Eq. (15) may be inaccurate for SG-DEXSY in heterogeneous systems because the characteristic *b*^1/3^ or *ρ*^2^ scaling of the non-Gaussian regime(s) is not accounted for. Applying the same principle, holding the sum of 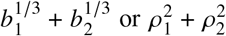 constant would potentially remove the effect of *f*_*NG,NG*_, or non-exchanging, non-Gaussian diffusion, but these two constancy conditions cannot be met simultaneously. In a follow-up work [81], we extended this curvature method to account for non-Gaussian diffusion by acquiring multiple *b*_*s*_ values to estimate *f*_*NG*_ and ⟨*c*_*NG*_⟩ prior to estimating *τ*_*k*_. Though the expression(s) became complicated, a key finding of that work is that non-Gaussian diffusion manifests itself as an *intercept* in the curvature that does not vary with *t*_*m*_. This finding suggests that while it may be difficult to measure *f*_exch_ in an absolute sense, the change in some signal quantity such as (*S*_end_ − *S*_mid_)/*S*_0_ w.r.t. *t*_*m*_ may be sufficient to characterize the exchange time *τ*_*k*_ via its *proportionality* with *f*_exch_. If said quantity is linear with *f*_exch_, even with some intercept, then *τ*_*k*_ can be measured robustly in a manner that is isolated from the effects of restriction or non-Gaussian diffusion.

### 3.4. Rapid quantification of exchange

Let us reconsider Eq. (13) for these points of interest: *S*_mid_/*S*_0_ along *ρ*_1_ = *ρ*_2_ ≈ 1.4, and *S*_end_/*S*_0_ along *ρ*_2_ ≈ 0 with *ρ*_1_ ≳ 1.4. Hereafter, we notate the equal *ρ* values in *S*_mid_/*S*_0_ as *ρ*_mid_ and the *ρ*_1_, *ρ*_2_ values for *S*_end_/*S*_0_ as (*ρ*_end,1_, *ρ*_end,2_). Note that *ρ*_end,2_ cannot be set to 0 exactly for SG measurements as the gradient is “always-on”. In the case of *S*_mid_, *ρ*_mid_ = 1.4 (*b* ≈ 2.3 ms/*μ*m^2^ for *D*_0_ = 2.15 *μ*m^2^/ms) should be large enough that signal in the Gaussian environment(s) during both *τ*_1_ or *τ*_2_ will be fully dephased — see again Fig. 2b — leaving only the terms in Eq. (13) with at least one encoding residing in the non-Gaussian environment(s):

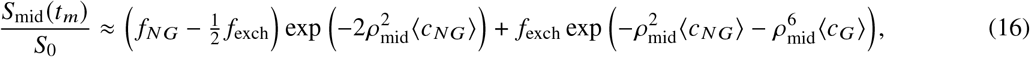

This expression is itself a linear relationship with *f*_exch_, with intercept 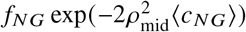 and a (negative) slope of exp 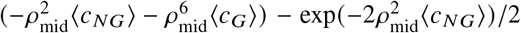. This linearity was hinted at in Figs. 4a and b, where the exchange contrast Δ*S*/*S*_0_ was seen to double as *f*_exch_ doubled. We can confirm this relationship by looking at simulation data, for which the position of walkers during each encoding can be tracked, i.e., the true *f*_exch_ is known.

Specifically, we define a walker as having exchanged if its position at the beginning of the simulation differs from that at the start of the second diffusion encoding period (in a binary sense: inside vs. outside of a sphere). If Eq. (16) holds, then fitting the exponential *decay* of *S*_mid_/*S*_0_ w.r.t. *t*_*m*_ should yield the same time-dependence (i.e., with *τ*_*k*_) as fitting the *growth* of *f*_exch_. Practically, this fit of *S*_mid_/*S*_0_ w.r.t. *t*_*m*_ needs at least 3 parameters without *a priori* knowledge. These parameters can be conceptualized as arising from (i) the decay of the equilibrium signal pools and any exchange during the encoding, which leads to an intercept at *t*_*m*_ = 0, (ii) a limit that is reached as *t*_*m*_ → ∞ and *f*_exch_ → *f*_exch, ss_, and (iii) a first-order exchange time, *τ*_*k*_. The fit has the general form *f* (*t*_*m*_) = *β*_1_ exp (− *β*_2_ *t*_*m*_) +*β*_3_ [23, 43], where *β*_2_ = 1/*τ*_*k*_ = *k*.

In Fig. 5, we plot the ground-truth *f*_exch_ and *S*_mid_/*S*_0_ vs. *t*_*m*_ for 3 simulated repetitions with *ρ*_mid_ ≈ 1.397 (*τ* = 0.59 ms) and *t*_*m*_ = [0.1, 0.5, 1, 2, 5, 10, 15, 20, 50] ms. Fits to each repetition yield *τ*_*k*_ ≈ 21.7 ± 0.9 and *τ*_*k*_ = 20 ± 5 ms (mean ± SD) for *f*_exch_ and *S*_mid_/*S*_0_, respectively. The exchange times thus agree between the curves, as predicted by Eq. (16). The estimation is also seen to be robust to a small amount of exchange during the encoding which is captured in the intercept *β*_3_. In principle, therefore, it is possible to measure *τ*_*k*_ from the decay of *S*_mid_/*S*_0_ along at least 3 points in *t*_*m*_ to fit the 3-parameter model, which is a highly efficient and quantitative measurement of exchange. Remarkably, this estimation can be performed without invoking any microstructural signal model and arises merely out of the signal decay of *S*_mid_/*S*_0_ itself, although we do assume that said decay takes a monoexponential form consistent with barrier-limited exchange.

**Figure 5:**
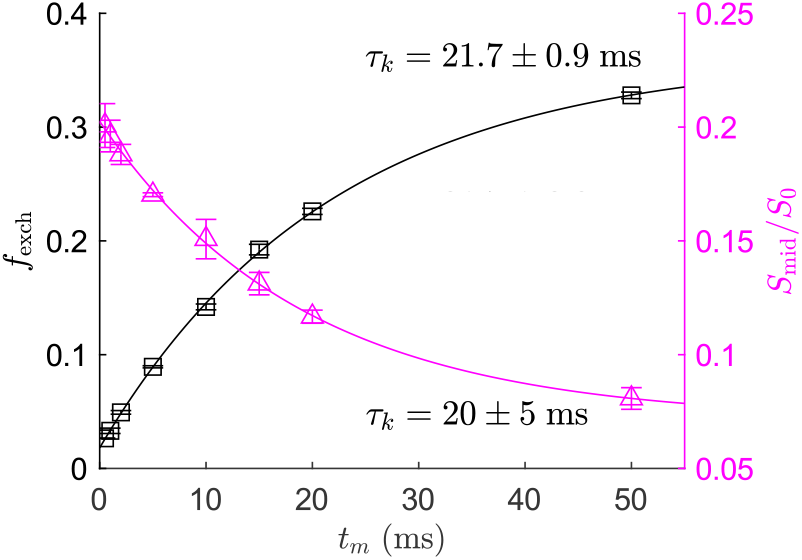
Comparison of *f*_exch_ and *S*_mid_/*S*_0_ obtained from simulation data with *ρ*_1_ = *ρ*_2_ ≈ 1.397 and *t*_*m*_ = [0.1, 0.5, 1, 2, 5, 10, 15, 20, 50] ms. Here, the ground-truth *f*_exch_ is quantified as the fraction of walkers that moved from inside/outside of a sphere between the start of the simulation and the start of the second diffusion encoding. The curve for *f*_exch_ (left axis, black) is fit to the form *f* (*t*_*m*_) = *β*_1_ [1 − exp (− *β*_2_ *t*_*m*_)] + *β*_3_ whereas *S*_mid_/*S*_0_ (right axis, magenta) is fit to *f* (*t*_*m*_) = *β*_1_ exp (− *β*_2_ *t*_*m*_) + *β*_3_. Note that a small intercept of *β*_3_ ≈ 0.02 is estimated in *f*_exch_ due to exchange during the encoding period. Error bars indicate mean ± SD from 3 repetitions. Solid lines are a fit to the mean. Fits to each repetition yield *τ*_*k*_ = 1/*β*_2_ = 21.7 ± 0.9 and 20 ± 5 for *f*_exch_ and *S*_mid_/*S*_0_, respectively. The values are in agreement, though noisier for *S*_mid_.

What of *S*_end_? For this point, we can again simplify Eq. (13) by assuming that signal which is in the Gaussian environment during the large first diffusion encoding has vanished:

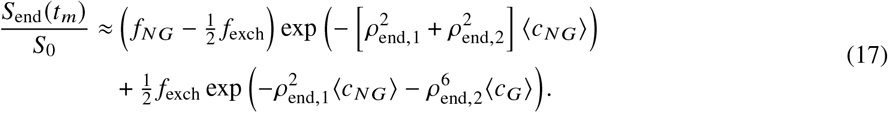

Similar to *S*_mid_/*S*_0_, this expression too can be described as a slope and intercept in *f*_exch_, though the slope is much smaller because *ρ*_end,2_ ≈ 0. Normalizing or subtracting *S*_mid_/*S*_0_ by a point such as *S*_end_/*S*_0_ (as in the curvature method) should thus have no effect on the fundamental linearity with *f*_exch_. The estimation of *τ*_*k*_ remains robust regardless. The choice of this additional point does become important if we consider the effect(s) of relaxation.

### 3.5. Accounting for relaxation

Thus far, we have ignored *T*_1_ relaxation during *t*_*m*_ by expressing the signals as normalized by *S*_0_. Normalizing for relaxation in the SG-DEXSY experiment is not straightforward, however. The *T*_1_ for the exchange-weighted point *S*_mid_ is not the ensemble *T*_1_ as measured by *S*_0_, but rather a *diffusion-weighted T*_1_ that is dominated by smaller compartments. Simply using *S*_mid_/*S*_0_ may leave some residual effect of *T*_1_ that biases the exchange measurement. The issues caused by *T*_1_ relaxation in these measurements was explored in detail by Williamson *et al*. [43] and approaches were given to normalize it. In general, these approaches exploit the fact that *S*_end_ is equivalently diffusion-weighted but is nominally *not* exchange-weighted (see again the rightmost “difference” column in Fig. 4). Therefore, we can use the decay of *S*_end_ w.r.t. *t*_*m*_ to characterize the diffusion-weighted *T*_1_ and remove it from *S*_mid_, recovering the linear relationship with *f*_exch_ in Eq. (16) that permits robust exchange measurement.

A straightforward approach is to take a ratio of the two points *S*_mid_/*S*_end_, i.e., normalizing by *S*_end_ rather than *S*_0_. One could also fit *S*_end_ separately before dividing out this decay from *S*_mid_. The latter approach has the benefit of requiring as few as 2 points in *S*_end_ while also avoiding noise propagation, which may be critical if SNR is low. Other approaches are also possible. Again, we defer to ref. [43] (where the ratio is called Method 2) for a more thorough comparison. Here, we choose the ratio approach for its simplicity and to avoid additional fitting steps.

We have thus arrived at the Diffusion Exchange Ratio (DEXR) method, which is comprised of the following fit:

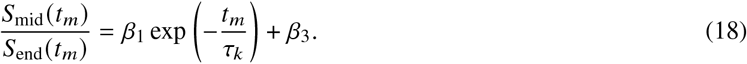

In Fig. 6a, we plot the raw signal values of *S*_mid_ and *S*_end_ acquired in a viable, *ex vivo* spinal cord with *ρ*_mid_ ≈ 1.4, *ρ*_end_ ≈ 1.56 (with a small *ρ*_2_ ≈ 0.81), and across 11 values of *t*_*m*_ = 0.2 − 300 ms (see caption). In terms of *b*-values and the curvature method, these parameters correspond to *b*_*s*_ = 4.5 and Δ*b*_*d*_ = 4.3 ms/*μ*m^2^. We see that both *S*_mid_ and *S*_end_ evolve by a diffusion-weighted *T*_1_ that is nearly identical at long *t*_*m*_, i.e., at steady state (see ref. [43] for estimates of *T*_1_ across many samples that confirm this), while exchange manifests as an additional decay in *S*_mid_. In Fig. 6b, we plot *S*_mid_/*S*_end_ along with a fit of Eq. (18) to the mean. The data takes roughly the expected form for a first-order exchange model (i.e., exponential decay to a baseline) after removing *T*_1_. Fits to each repetition yield *τ*_*k*_ = 11 ± 3 ms, which is consistent with our previous reports [23, 43, 63].

**Figure 6:**
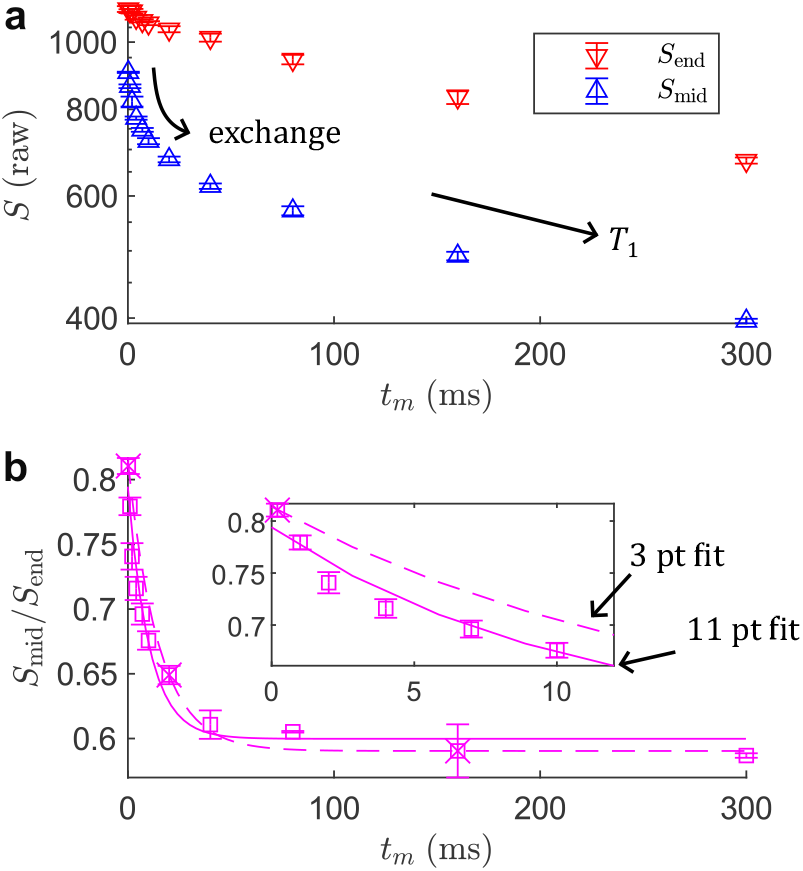
Estimation of the exchange time *τ*_*k*_ from two points per *t*_*m*_ (*S*_mid_ and *S*_end_) in viable *ex vivo* spinal cord. The normalization and fitting approach shown here comprise the DEXR method. **(a)** Plots of the raw signal decay of *S*_mid_ with *ρ*_mid_ ≈ 1.39 (*τ* ≈ 0.59 ms) and *S*_end_ with *ρ*_end_ ≈ 1.55 (*τ* ≈ 0.74 ms) for 11 values of *t*_*m*_ = [0.2, 1, 2, 4, 7, 10, 20, 40, 80, 160, 300] ms. Data is plotted on a log *y*-axis to highlight the approximately linear decay at long *t*_*m*_, indicative of diffusion-weighted *T*_1_ relaxation. Error bars indicate mean ± SD from 3 repetitions on the same sample. Both points evolve by *T*_1_, while *S*_mid_ also evolves due to exchange. Fitting a monoexponential decay to *S*_end_ yields an apparent diffusion-weighted *T*_1_ ≈ 600 ± 20 ms, which differs from the *T*_1_ ≈ 710 ± 10 ms obtained by fitting an *S*_0_ acquisition with *τ*_1_ = *τ*_2_ = 0.05 ms (fits and data not shown, see ref. [43]), highlighting the non-triviality of accounting for *T*_1_. Note that because *S*_end_ is not acquired precisely at *τ*_2_ = 0, but at *ρ*_2_ ≈ 0.81, this point is also slightly exchange-weighted — see the non-linear behavior at short times. **(b)** Fit of Eq. (18) to the ratio *S*_mid_/*S*_end_, yielding *τ*_*k*_ = 11 ± 3 ms. Fits to the mean using all 11 *t*_*m*_ (solid line) or a minimal 3 values of *t*_*m*_ = [0.2, 20, 160] ms (crosses, dashed line) are plotted. The minimal sampling yields a similar *τ*_*k*_ = 17 ± 4 ms.

To demonstrate the potential efficiency of the method, we perform the same fit using a minimal 3 values of *t*_*m*_ = [0.2, 20, 160] ms (indicated by crosses and a dashed line in Fig. 6b). Using 3 points along *t*_*m*_ or 6 points in total yields *τ*_*k*_ = 17 ± 4 ms. Thus, similar parameters and variation can be obtained from minimal data, though a slightly smaller *τ*_*k*_ is estimated using the full dataset. This may be due to some multiexponential character in the data, which can be seen in the zoomed inset in Fig. 6b. The behavior is interesting and may indicate that a first-order exchange model is insufficient to explain the data, which we will explore further in the following section on time-dependent diffusion (see Ordinola *et al*. [89] and Cai *et al*. [90] for other investigations of this phenomenon).

It should be mentioned that there are other effects in the DEXR experiment. For instance, there is also a small difference in *T*_2_-weighting between *S*_mid_ and *S*_end_ due to their different *τ* values, as well as the possibility of *T*_2_-*T*_2_ exchange, though we expect that these effects will be small given that *τ* < 1 ms ≪ *T*_2_. Another issue is that in our SG-DEXSY implementation, *S*_end_ is slightly exchange-weighted (see Methods and Fig. 6a) and dividing it removes some exchange contrast [43]. Nonetheless, these effects will not impact *τ*_*k*_ estimates much because they are captured in the other fit parameters *β*_1_ and *β*_3_ that characterize the range of signal variation. We reiterate that the linearity between the ratio *S*_mid_/*S*_end_ and *f*_exch_ is what is important and this is preserved and robust to confounding effects. That said, Eq. (18) and its demonstration in Figs. 5 and 6 form the basis of the DEXR method.

### 3.6. Extracting restriction parameters

Although *τ*_*k*_ is the main parameter of interest, *β*_1_ and *β*_3_ may also hold important information about exchanging pools and their environment. If the confounding effects such as exchange during the encoding can be accounted for, then these parameters contain information about the restricting microenvironment and can potentially be used to estimate *f*_*NG*_ and ⟨*c*_*NG*_⟩. Consider that the total signal variation *β*_1_ should be related to *f*_exch, ss_, with a larger *β*_1_ indicating larger *f*_exch, ss_, all else being equal. The intercept where *t*_*m*_ = 0, given by *β*_1_ + *β*_3_, should be related to the decay of the equilibrium signal fractions as well as exchange during the encoding. Can these terms be rearranged to yield restriction parameters?

First, let us try to estimate *f*_*NG*_. Consider that by taking some ratio in combinations of *β*_1_ and *β*_3_, we can remove any leading exponential attenuation terms. We will leave aside the issue of *S*_end_ being slightly exchange weighted for now, working with an idealized *S*_mid_/*S*_0_ from Eq. (16). The limiting behavior(s) can be written following some rearrangement as:

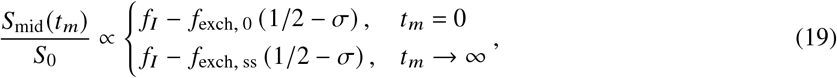

where we leave out the leading decay term exp 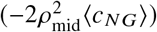 for compactness, and where

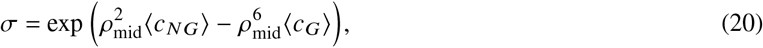

can be thought of as a filter efficiency that characterizes how well a single encoding with *ρ*_mid_ separates the Gaussian and non-Gaussian signal, and *f*_exch, 0_ is the exchange that transpires during the first encoding. More specifically, *σ* describes the degree to which signal that has exchanged (i.e., which spends one of the two encodings in the Gaussian environment) is dephased relative to the non-exchanging, non-Gaussian signal. An appreciable value of *σ* indicates that there remains some coherent exchanged signal that contributes to *S*_mid_ such that the second term in Eq. (16) cannot be ignored. Taking the ratio of the total signal variation and the intercept, *β*_1_/(*β*_1_ + *β*_3_), we obtain

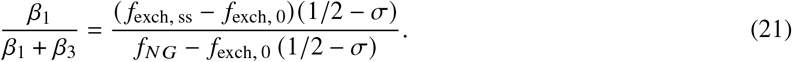

Substituting *f*_exch, ss_ = 2 *f*_*NG*_ (1 − *f*_*NG*_) and dividing *f*_*NG*_,

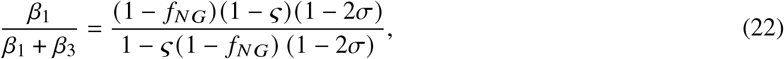

where

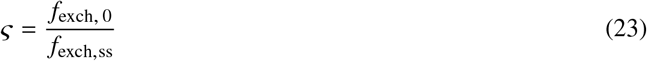

captures how much of the total exchange is missed in the first encoding, and we note that *f*_exch, 0_ = 2*ς f*_*NG*_ (1 − *f*_*NG*_). Rearranging,

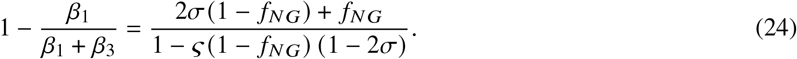

We see that if the confounding effects can be ignored — i.e., if both *σ, ς* = 0 — then the right-hand-side is simply *f*_*NG*_. Thus, *f*_*NG*_ can potentially be experimentally measured from the same data and fit, with the following simplifying cases:

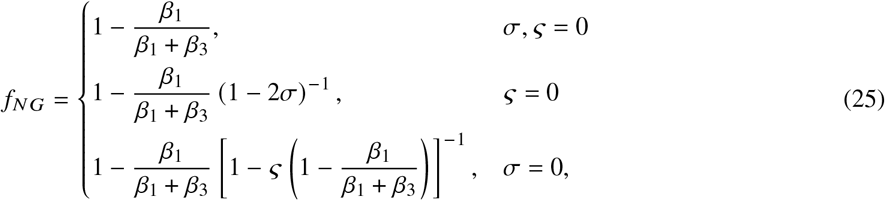

where the *σ, ς* = 0 case is readily extracted from DEXR data and the other cases describe possible corrections.

Practically, *σ* reduces the slope between *f*_*NG*_ and the quantity 1 –*β*_1_/(*β*_1_/*β*_3_)such that its effect is to bias *f*_*NG*_ upwards when compared to the *σ, ς* = 0 case. The effect is more pronounced for smaller *f*_*NG*_. The effect of *ς* is similar in that it also biases *f*_*NG*_ upwards compared to the *σ, ς* = 0 case. Recall that the relationship between *f*_exch, ss_ and *f*_*NG*_ is quadratic, see Eq. (14); therefore *ς* will introduce an upwards bowing in *f*_*NG*_ vs. 1 − *β*_1_/(*β*_1_ +*β*_3_), with the maximal effect at *f*_*NG*_ = 0.5. In Fig. 7, we plot *f*_*NG*_ vs. 1 − *β*_1_/(*β*_1_+*β*_3_) for various values of *σ* and *ς*. The curves indicate that *σ* can have a large effect on *f*_*NG*_ estimates, while the effect of *ς* is comparatively small. This suggests that when selecting *ρ*_mid_, it is preferable to err on the side of larger *ρ* in order to better crush the Gaussian signal and yield robust *f*_*NG*_ estimates. Given that the optimal range in Fig. 3 is quite broad, this should have little effect on the SNR of *τ*_*k*_ estimates. In all cases, the effect is roughly linear such that we can correct *f*_*NG*_ reasonably well simply by drawing a line between *f*_*NG*_ = 0 and *f*_*NG*_ = 1. From Eq. (24) we obtain:

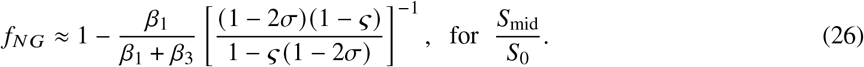

Note that *σ, ς* can never actually be 0 and the bracketed term above is always > 1 (inverse < 1). As such, using 1 − *β*_1_/(*β*_1_ + *β*_3_) as an estimate of *f*_*NG*_ is a systematic overestimation, the size of which roughly scales with 1 − *f*_*NG*_ = *f*_*G*_.

**Figure 7:**
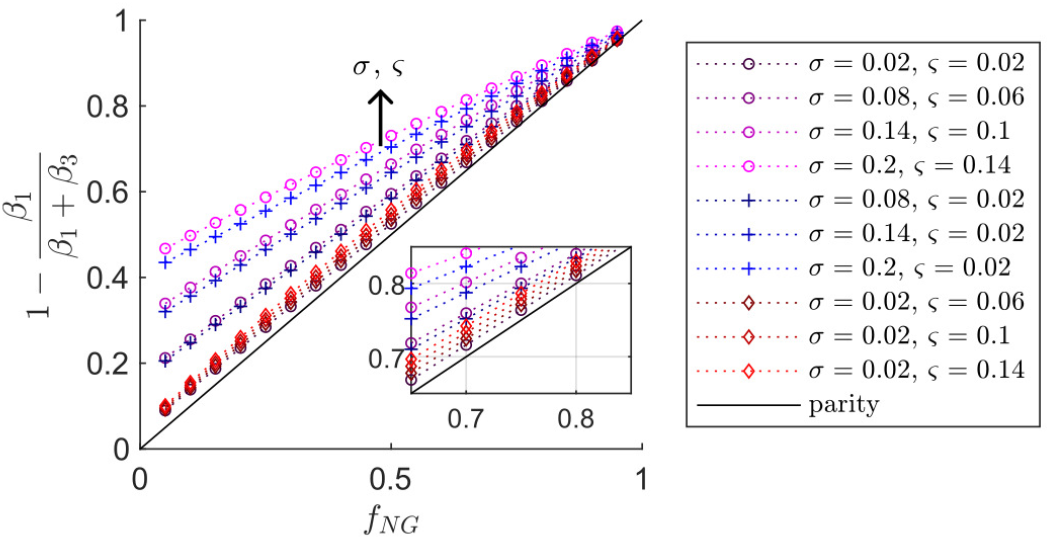
Relationship between *f*_*NG*_ and the fit-derived quantity 1 − *β*_1_/(*β*_1_ + *β*_3_) for various values of 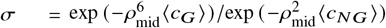 and *ς* = *f*_exch, 0_/*f*_exch,ss_, which characterize the confounding effects of extant exchanged signal and exchange during the encoding, respectively. The solid black line indicates parity when *σ, ς* = 0. Curves of Eq. (24) derived from an idealized *S*_mid_/*S*_0_ are plotted for *σ* = [0.02, 0.08, 0.14, 0.2] and *ς* = [0.02, 0.06, 0.1, 0.14]. The parameters are varied together (magenta circles) and independently (blue crosses, red diamonds), with deepening color representing increasing values. In all cases, the behavior manifests, roughly speaking, as a decrease in the linear relationship or slope between *f*_*NG*_ and *β*_1_/(*β*_1_ + *β*_3_).

Another confounding effect arises from the exchange weighting in *S*_end_. As mentioned in Eq. (17), an idealized *S*_end_/*S*_0_ also has a slope and intercept in *f*_exch_ if *ρ*_end,2_ > 0. Giving a similar treatment to *S*_end_/*S*_0_ from Eq. (17) as in Eq. (19), we obtain

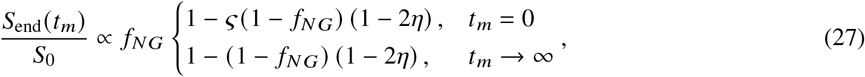

which is similar to Eq. (19) but with *η* instead of *σ*, and where

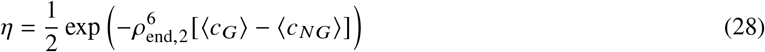

is a term that characterizes the decay of exchanged signal due to *ρ*_end,2_, which we have approximated as being Gaussian for all environments since *ρ*_end,2_ < 1. Again, we leave out the leading attenuation term 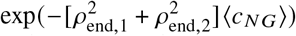 for compactness. As expected, if *ρ*_end,2_ = 0, then *η* = 1/2 and *S*_end_/*S*_0_ has no *t*_*m*_ dependence. Note that *ς* actually differs between *S*_end_ and *S*_mid_ because their value(s) of *τ* differ. Practically, this difference in *τ* is small ≈ 0.14 ms due to the high *g* used here, and we will assume that *ς* is approximately equal in both points. Furthermore, the effect of *ς* in Fig. 7 is small such that this approximation should not affect the *f*_*NG*_ estimate significantly. If we can assume that *ς* is the same, then we can simply “add back” the *t*_*m*_ dependence that is lost by dividing *S*_end_, replacing (1 − 2*σ*) with (1 − 2*σ*) + (1 − 2*η*) = 2(1 − *σ* − *η*) wherever it appears. Thus, we approximate from Eq. (26) that

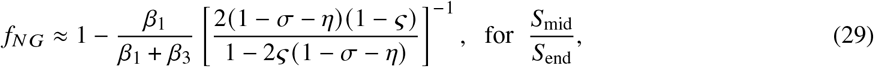

taking into account all three effects or corrections from *σ, ς*, and *η*: incomplete dephasing of exchanged signal, exchange during the first encoding, and exchange weighting in *S*_end_, respectively. Importantly, the general linear behavior with intersection at *f*_*NG*_ = 1 shown in Fig. 7 is preserved in Eq. (29).

Let us assess expected values of *σ, ς*, and *η*. The value of *ς* can be estimated from the fit itself — using *τ*_*k*_ ≈ 11 ms and 2*τ* ∼ 1 ms, we obtain *ς* ≈ 0.1. However, *σ* and *η* cannot be estimated from the data alone. Using the values ⟨*c*_*NG*_⟩ = 0.18, ⟨*c*_*G*_⟩ = 0.26 obtained for spinal cord in Fig. 2, we estimate that *σ* could be as high as ≈ 0.2 for *ρ*_mid_ = 1.4, though we again stress that the SG-SE fits are suspect to aforementioned confounds, and ⟨*c*_*G*_⟩ is likely underestimated. As an upper-bound, using the maximal ⟨*c*_*G*_⟩= 2/3 corresponding to free Gaussian diffusion yields just *σ* ≈ 0.01 using the same ⟨*c*_*NG*_⟩. A lower-bound can be estimated from literature values of the tortuosity of the ECS

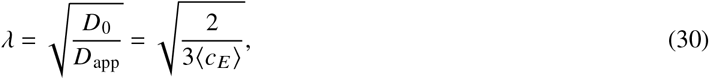

which generally fall below *λ* ≈ 1.7 (and may be much smaller in neonatal mouse tissue that has larger ECS occupancy compared to adult tissue) [91]. Using *λ* ≲ 1.7 gives ⟨*c*_*G*_⟩≳ 0.5, yielding *σ* ≲ 0.04. For *ρ*_end,1_ = 1.55, *ρ*_end,2_ = 0.81, we obtain 0.44 ≲ *η* ≤ 0.47. To a first approximation, we estimate that the bracketed correction term in Eq. (29) may range from ≈ 0.9 − 1.02 in spinal cord data. Surprisingly, these effects when considered together yield a correction close to 1. Thus, *f*_*NG*_ = 1 − *β*_1_/(*β*_1_ + *β*_3_) may be a good estimate in this data, particularly for larger values of 1 − *β*_1_/(*β*_1_ + *β*_3_) > 0.7.

Let us now isolate ⟨*c*_*NG*_⟩. Of course, the estimations of *f*_*NG*_ and ⟨*c*_*NG*_⟩ are actually coupled via the various correction terms and the two cannot be truly isolated. Nonetheless, we can proceed with estimating some apparent ⟨*c*_*NG*_⟩ by assuming that our initial *f*_*NG*_ estimate is accurate. We have at *t*_*m*_ → ∞ for *S*_mid_/*S*_0_ that

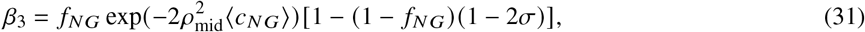

which removes *ς*. Thus,

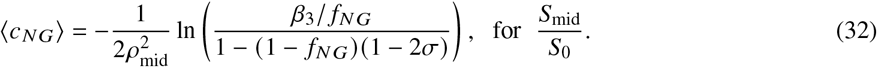

And similarly,

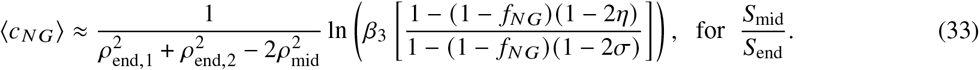

Note that if both restriction and exchange are quantified, then we can go further and calculate the effective permeability from Eqs. (9) and (10) as a secondary result,

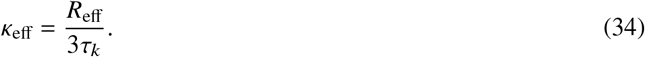

where 3/*R*_eff_ is the SVR of the effective sphere, though this can easily be adapted for other geometries simply by changing the geometric prefactor.

### 3.7. Restriction results

Let us revisit the results in Figs. 5 and 6 and estimate *f*_*NG*_ and ⟨*c*_*NG*_⟩. In the simulation data (Fig. 5), we obtain *f*_*NG*_ = 0.34 ± 0.06 from Eq. (26) assuming that *σ, ς* = 0 and ⟨*c*_*NG*_⟩ = 0.17 ± 0.07, *R*_eff_ = 0.88 ± 0.08 *μ*m using the expressions for *S*_mid_/*S*_0_ in Eqs. (26) and (32). These parameters agree well with the ground truth of *R* = 0.95 *μ*m and *f*_*NG*_ ≈ 0.34, though *R* is slightly underestimated due perhaps to a truncation of Neuman’s [5] expressions to arrive at Eq. (4). What of the correction terms? From the *f*_exch_ fit in Fig. 5, we estimate that *ς* ≈ 0.04. Again, *σ* cannot be determined from the data itself because of the lack of sensitivity to ⟨*c*_*G*_⟩, but we point out that a value close to ⟨*c*_*G*_⟩ = 2/3 is reasonable given the loose packing of these spheres [92]. For the mean ⟨*c*_*NG*_⟩ above and a somewhat arbitrary ⟨*c*_*G*_⟩ = 0.6, we have *σ* ≈ 0.015. From Eq. (29), we obtain a slightly smaller *f*_*NG*_ = 0.33 ± 0.06 and *R*_eff_ = 0.84 ± 0.1 *μ*m. Note that applying these corrections updates the estimated ⟨*c*_*NG*_⟩ and thereby the correction terms themselves. We could perform the correction iteratively until the parameters converge, but because ⟨*c*_*G*_⟩ has the greatest effect on *σ* and *η*, this is not necessary and one iteration suffices. The permeability estimated from *τ*_*k*_ and the corrected *R*_eff_ using Eqs. (34) and (9) is *κ*_eff_ = 0.14 ± 0.05 *μ*m/ms, which can be compared to a ground-truth estimate from the *f*_exch_ curve in Fig. 5 and *R* = 0.95 *μ*m, which yields *κ*_eff_ = 0.094 ± 0.007 *μ*m/ms. Note that an underestimation of *R*_eff_ will lead to a corresponding overestimation in *κ*_eff_ according to Eq. (34).

These parameters agree more closely with the ground truth than the fit of Eq. (7) to simulated SG-SE data, shown in Fig. 2, particularly for *f*_*NG*_. In that fit, *f*_*NG*_ ≈ 0.44 was overestimated. Consider that in SG-DEXSY, walkers have more time over *t*_*m*_ to explore the tortuous space and manifest as hindered signal, rather than appearing as restricted over the short timescale of an SG-SE. We reiterate that in DEXR data, the estimation of exchange and restriction are isolated, with exchange being the only effect that influences the time-dependence with *t*_*m*_, while the effect(s) of restriction are estimated using only the other fit parameters that capture the initial and limiting behavior of the signal (i.e., *β*_1_ and *β*_3_, along with the various corrections).

For the fully sampled spinal cord data in Fig. 6, we obtain *f*_*NG*_ = 0.752 ± 0.003, ⟨*c*_*NG*_⟩ ≈ 0.28 ± 0.02, and *R*_eff_ = 1.07 ± 0.02 *μ*m, without correction. With this mean ⟨*c*_*NG*_⟩, we estimate the correction terms using a lower-bound ⟨*c*_*G*_⟩ = 0.5 corresponding to *λ* ≈ 1.7, yielding *σ* ≈ 0.04 and *η* ≈ 0.47. With correction: *f*_*I*_ = 0.746 ± 0.003, *R*_eff_ = 1.11 ± 0.02 *μ*m, and *κ*_eff_ = 0.33 ± 0.09 *μ*m/ms. This value of *κ*_eff_ is large, but is within the range of permeability values expected for phospholipid bilayers that highly express aquaporin water channels [93, 94], such as those found in GM. Solenov *et al*., for example, report *κ* ≈ 0.5 *μ*m/ms in primary cultures of mouse astrocytes, measured via calcein fluorescence quenching [95].

As was the case for the simulation data shown in Fig. 5, a different *f*_*NG*_ ≈ 0.75 is obtained here than in Fig. 2, where a much smaller *f*_*NG*_ ≈ 0.16 was estimated for spinal cord, though that sample was fixed. Given the very fast exchange time of *τ*_*k*_ ≈ 11 ms, the confounding effect of exchange during the SG-SE encoding may have been significant, potentially leading to a decreased *f*_*NG*_. While the ground truth in this case is unknown (as are the effects of fixation, which permeabilizes membranes [96]), consider that *f*_*NG*_ ≈ 0.75 estimated using DEXR roughly agrees with the expected occupancy fraction of the ICS *in vivo*; the ECS is reported as occupying between ∼ 15 − 30% of the space in rodent spinal cord, specifically [91], in accordance with *f*_*G*_ = 1 − *f*_*NG*_ ≈ 0.25. We speculate that DEXR is more quantitatively accurate than SDE, highlighting once again the key advantage of the method in isolating exchange from restriction. That said, signal from water in the tissue ICS and ECS would not be expected to exactly parse into *f*_*NG*_ and *f*_*G*_; heterogeneity of plasma membrane length scales may lead to some water in the ICS appearing as more mobile or unrestricted, and the narrow width of the ECS may lead to some water in the ECS appearing as restricted. We cannot know, truly, what *f*_*NG*_ is within tissue, though the obtained estimate is reasonable.

To summarize, the fit parameters obtained from simulation and spinal cord DEXR data are provided in Table 1. We highlight again that the simulation data produces accurate estimates of *τ*_*k*_ and *f*_*NG*_ compared to the ground truth, and reasonable estimates of *R*_eff_ and *κ*_eff_, though some systematic over/underestimation remains. In the spinal cord data, we also explored the feasibility of data reduction, and compared 11 vs. 3 mixing times (Fig. 6b). The use of just 3 mixing times is a vast reduction in data requirement compared to conventional DEXSY.

**Table 1:** Exchange and restriction parameters estimated using DEXR data from simulation and from viable, *ex vivo* neonatal mouse spinal cord. For simulation data, *ρ*_mid_ ≈ 1.397 (*S*_end_ was not simulated). For spinal cord, *ρ*_mid_ ≈ 1.4 and (*ρ*_end,1_, *ρ*_end,2_) ≈ (1.56, 0.81). Error bars = mean ± SD from 3 repetitions on the same sample. In all data sets, Eq. (18) was first fit to yield *τ*_*k*_, *β*_1_, and *β*_3_. Subsequently Eqs. (26) and (32) were evaluated for simulation data, and Eqs. (29) and (33) were evaluated for spinal cord to yield *f*_*NG*_ and *R*_eff_ from Eq. (9). See the main text for correction terms (*ς, σ, η*). Subsequently, Eq. (34) was used to yield *κ*_eff_. The estimated parameters for simulation data can be compared to the simulation ground truth shown in Fig. 5.

| Data source | Sampling in $t_m$ | $\tau_k$ (ms) | $f_{NG}$ | $R_{\text{eff}}$ ($\mu\text{m}$) | $\kappa_{\text{eff}}$ ($\mu\text{m}/\text{ms}$) |
| --- | --- | --- | --- | --- | --- |
| simulation (ground truth) | 9 pt., 0.1 – 50 ms | $21.7 \pm 0.9$ | 0.337 | 0.95 | $0.094 \pm 0.007$ |
| simulation | 9 pt., 0.1 – 50 ms | $20 \pm 5$ | $0.33 \pm 0.06$ | $0.84 \pm 0.1$ | $0.14 \pm 0.05$ |
| viable spinal cord | 11 pt., 0.2 – 300 ms | $11 \pm 3$ | $0.746 \pm 0.003$ | $1.11 \pm 0.02$ | $0.33 \pm 0.09$ |
| viable spinal cord | 3 pt., [0.2, 20, 160] ms | $17 \pm 4$ | $0.71 \pm 0.01$ | $1.10 \pm 0.01$ | $0.21 \pm 0.06$ |

## 4. Alternative analysis with time-dependent diffusion

Having provided a pipeline to analyze DEXR data to yield both exchange (*τ*_*k*_, *κ*_eff_) and restriction parameters (*f*_*NG*_, *R*_eff_) we now turn towards an alternative analysis in terms of time-dependent diffusion and show that the same data can be used to yield an apparent VACF.

### 4.1. A time-domain signal representation

The conventional measurement of time-dependent diffusion using TDS is based on a frequency-domain representation of the signal [45, 46, 48, 49]

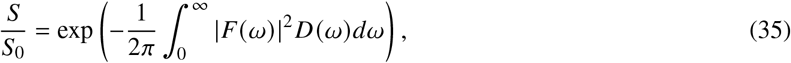

where *F* (*ω*) is the (truncated) spectrum of *F* (*t*), where 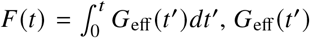 is the effective gradient, and *D*(*ω*) is the spectrum of the 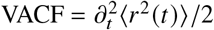. To be more explicit about how these different transport quantities (in a single dimension) are related, the conversions between them are summarized as:

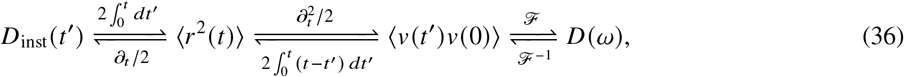

where *ℱ* denotes a Fourier transform, *ℱ*^− 1^ its inverse, and with the additional relation *D* (*t*) = ⟨*r*^2^ (*t*)⟩/2*t*. We see that *D*_inst_ *t* is half the first derivative of the MSD w.r.t. time, while the VACF is half the second derivative. The frequency-domain expression in Eq. (35) is useful in the case of gradient sequences with a sharp power spectrum, but less useful in describing the time- or frequency-dependence of more general diffusion MR sequences. According to Ning *et al*. [55], Eq. (35) can be rewritten in several equivalent, time-domain representations. One of these expresses the signal in terms of the instantaneous diffusivity *D*_inst_(*t*) and the cumulative gradient autocorrelation function 𝒞 (*t*):

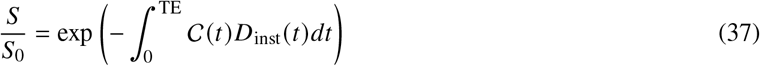

where 𝒞 (*t*) is given by

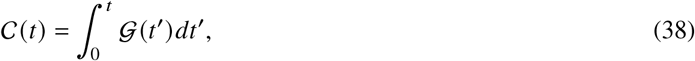

where

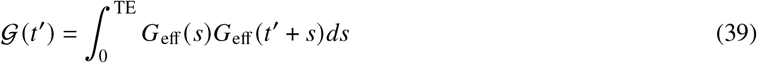

is the autocorrelation function of the effective gradient and TE is the time of echo formation. It is important to note that the *b*-value:

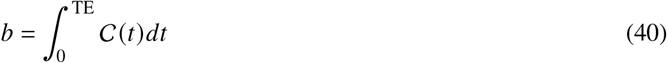

can be viewed in this representation as a multiple time integral of the autocorrelation function of the effective gradient waveforms. This explicitly opens up the possibility of using unconventional gradient waveforms as a means of refining the diffusion weighting, and reinforces the notion that the *b*-value sensitizes the signal to motional correlations between different encoding periods.

Using this signal representation in 𝒞(*t*), we can characterize the sensitivity of our SG-DEXSY sub-sampling scheme in the time domain. In the case of an *S*_end_ acquisition or simply an SG-SE experiment, we have that

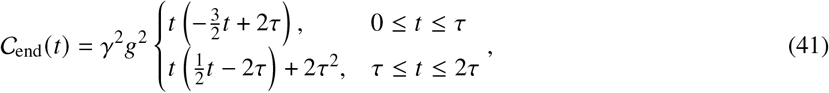

where *τ* = *τ*_1_, and assuming that *G*_eff_ ∈ {0, − *γg*, +*γg*} = 0 for *t* > 2*τ*_1_, i.e., we ignore the diffusion-weighting of the CPMG readout, see Fig. 1. As an aside, we take this opportunity to point out that there is a typo in Eq. (14) of Cai *et al*. [56], where a factor of 2 is missing in the second interval from *τ* < *t* ≤ 2*τ*. This 𝒞_end_ (*t*) is a single broad “lobe” centered at *t* = 2*τ*/3, with coarse sensitivity in the time-domain as would be expected of a non-oscillating sequence. For *S*_mid_ with *τ* = *τ*_1_ = *τ*_2_, the analogous expression for the final echo formed at TE = 4*τ*+*t*_*m*_ is tedious but straightforward to calculate:

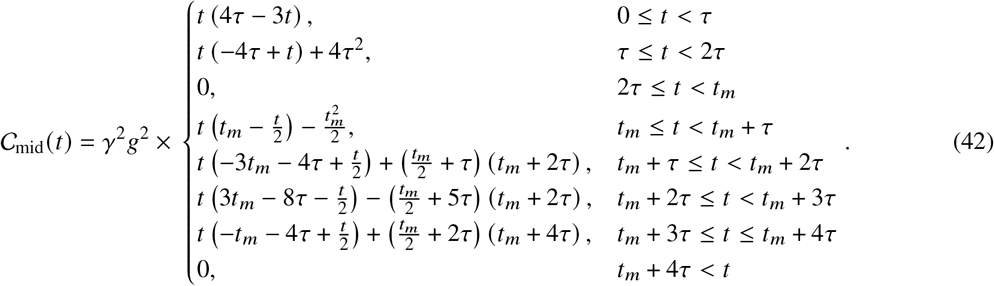

In Fig. 8, we plot 𝒞_end_ (*t*) and 𝒞_mid_ (*t*) for exemplar timing parameters consistent with the curvature method using *b*_*s*_ ≈ 4.5 ms/*μ*m^2^ in order to illustrate the shape of these time-domain weightings. The weighting over the timescale of the first encoding in either case is similar — indeed, these “lobes” integrate to the same total *b*-value of *b*_*s*_, though 𝒞_end_ (*t*) spans a wider time range. The 𝒞_mid_ (*t*) curve, however, has two additional lobes centered about *t* = 2*τ*+*t*_*m*_, with the negative lobe having a peak at *t* = 4*τ*/3+ *t*_*m*_ and the positive lobe peaking at *t* = 8*τ*/3 +*t*_*m*_. These lobes arise from the autocorrelation between the first and second encodings and integrate to ∓*b*_*s*_/2. If we assume that the variation in *D*_inst_ (*t*) is small on the timescale of *τ* such that we can treat it as being approximately constant over each lobe, then

**Figure 8:**
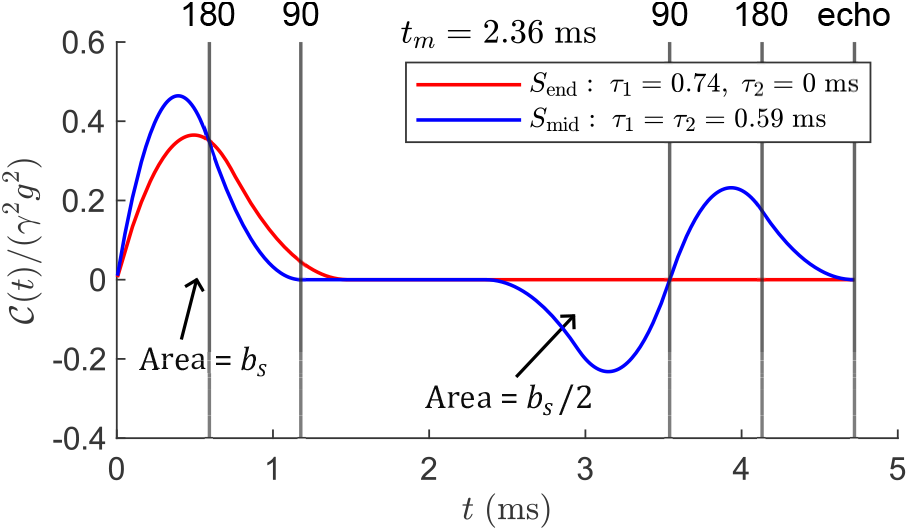
Time-domain weighting 𝒞(*t*) for an *S*_end_ acquisition with *τ*_1_ = 0.74 ms compared to an *S*_mid_ acquisition with *τ*_1_ = *τ*_2_ = 0.59 ms; *t*_*m*_ = 2.36 ms for both. These parameters are approximately consistent with the curvature method and *b*_*s*_ = 4.5 ms/*μ*m^2^. The curves are plotted on a non-dimensionalized *y*-axis of 𝒞(*t*)/(*γ*^2^*g*^2^), where *g* = 15.3 T/m. Gray vertical lines indicate the timing of RF pulses and echo formation for the *S*_mid_ acquisition. The weightings over 0 ≤ *t* < 2*τ* are similar between the two acquisitions, as expected for an equal total *b*-value or *b*_*s*_. However, 𝒞_mid_ *t* has an additional two lobes centered about *t* = 2*τ*+*t*_*m*_ with peaks at *t* = (4/3) *τ* +*t*_*m*_ and (8/3) *τ*+ *t*_*m*_. These peaks arise from the autocorrelation of the separated diffusion encodings and integrate to ∓*b*_*s*_/2, respectively, while the first lobe integrates to +*b*_*s*_ (i.e., if the *y*-axis were multiplied by *γ*^2^*g*^2^).

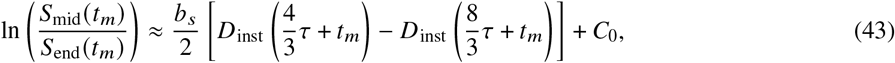

where *τ* here corresponds to *τ*_1_ = *τ*_2_ of *S*_mid_ and *C*_0_ represents a unitless, negative constant that accounts for any remaining contribution from the imperfect cancellation of the initial lobes in 𝒞_end_(*t*) and 𝒞_mid_(*t*). By dividing the effective spacing between the pair of positive and negative lobes, 4*τ*/3, this becomes a forward, first-order finite difference approximation of the slope in *D*_inst_(*t*) at *t* = 2*τ* + *t*_*m*_. Thus we can rearrange the above into an expression that is an experimental measurement of *∂*_*t*_ *D*_inst_(*t* = 2*τ* + *t*_*m*_), which is equivalently the VACF as shown in Eq. (36):

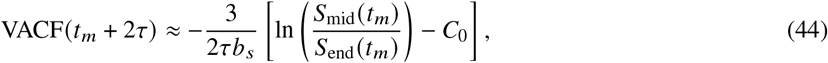

where we notate the VACF = ⟨*ν* (*t*)*ν* (0)⟩ as a function of time. Additionally, consider that at long times, we can approximate lim_*t* → ∞_ *∂*_*t*_ *D*_inst_ (*t*) ≈ 0 (i.e., the long-time behavior where *D*(*t*)≃*D*_app_ is reached, and the bracketed term in Eq. (43) vanishes). Therefore, *C*_0_ can potentially be approximated by the limiting value of *S*_mid_/*S*_end_, called *β*_3_ in the previous section:

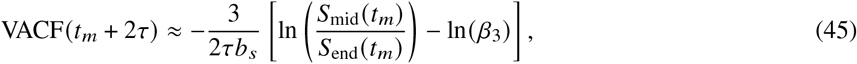

where

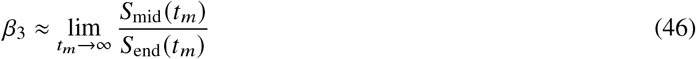

is estimated from a fit of Eq. (18). Such an approximation is justified by the data itself, as decay toward a baseline is clearly observed (see Figs. 5 and 6b). This baseline is when the VACF ≈ 0.

From this perspective, our method can be interpreted as a measurement of time-dependent diffusion — indeed, it is a *direct* measurement of the VACF — wherein the weighting in the time-domain is varied via *t*_*m*_. The effective resolution (i.e., the width of the positive and negative lobes), is the value of 2*τ* for *S*_mid_. This method can probe the VACF from *t* ≳ 2*τ* ≈ 1 ms to an upper limit depending on the SNR constraint imposed by the sample *T*_1_, which for this field strength is on the order of ∼ 1 s (see Fig. 6a), and by the chosen diffusion weighting. Using just one method, we can probe multiple orders of magnitude in the time domain.

### 4.2. The Gaussian phase approximation and stationarity

Before going further and applying Eq. (45) to the DEXR data presented previously, we stress that these time-dependent signal representations are valid if and only if the transport process is stationary with no net flow into/out of the active region, nor any re-partitioning of the signal between compartments (i.e., detailed balance). Another assumption is that the distribution of spin phases *P* (*ϕ*) is well-approximated by a Gaussian. If so, the first two cumulants suffice to describe *P* (*ϕ*). This is known as the Gaussian phase approximation (GPA), used since the infancy of diffusion MR [5, 57, 69, 97, 98]. The GPA holds in the motional averaging and Gaussian diffusion regimes but not in the localization regime. It holds in the motional averaging regime because the averaging process within a given restricted volume implies that each spin isochromat is in effect a random sample of the underlying *P* (*ϕ*) and the central limit theorem applies [69]. The GPA can be equivalently stated as there being negligible localized signal.

Recall that in our data, we have *ρ* ≳ 1, but no greater than *ρ* ≈ 1.6, and thus a significant amount of localized signal is not expected because *𝓁*_*d*_ and *𝓁*_*g*_ remain similar, and the GPA should hold. This can be explored in the simulation data. In Figs. 9a and b we show the phase distributions *P* (*ϕ*) at the end of the first and second encodings of the simulated SG-DEXSY experiment, respectively (i.e., at the times of echo formation). We see that the non-Gaussian, non-exchanging fraction *f*_*NG,NG*_ at the second echo is well-described by a Gaussian (*p* > 0.3, see caption), while the other signal fractions are nearly completely dephased. Therefore, the GPA holds overall. Given the similar *R*_eff_ (Table 1) estimated for spinal cord, Ning *et al*.’s [55] signal representations can be said to hold in the spinal cord data. The distributions shown in Fig. 9 also serve as a visual summary of the SG-DEXSY experiment for *S*_mid_, illustrating how the signal pools *f*_*G*_ and *f*_*NG*_ evolve over both encodings, and how the signal from *f*_exch_ is largely dephased, leading to proportionality between the ensemble signal and *f*_exch_.

**Figure 9:**
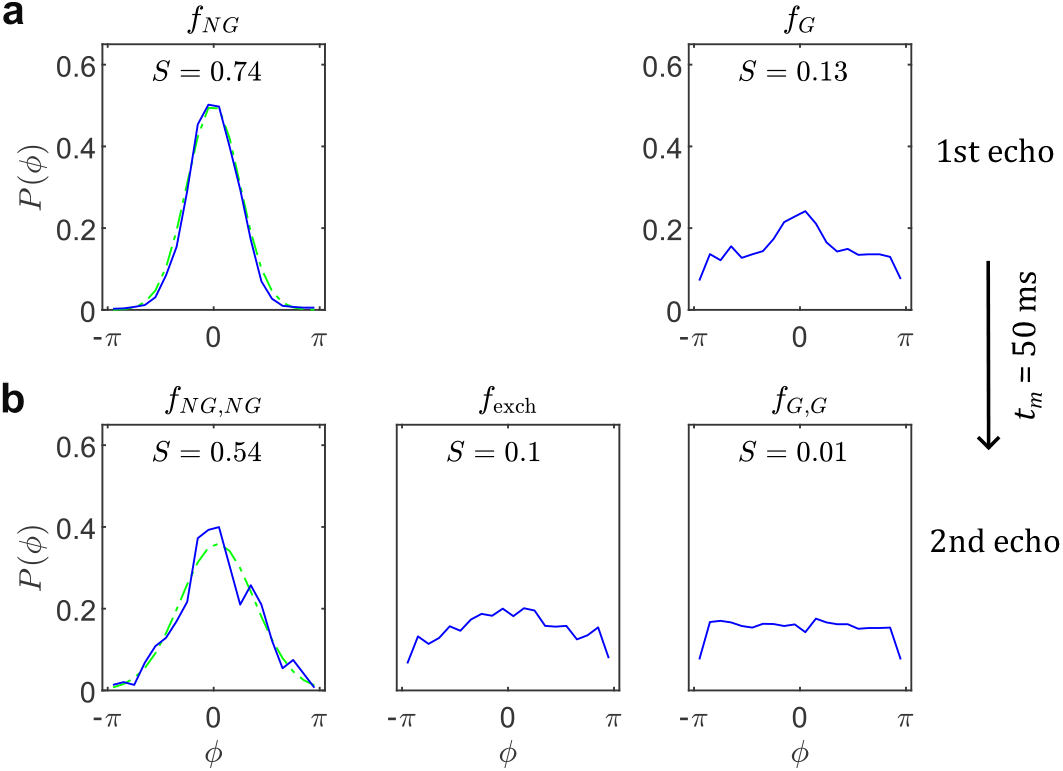
Phase distributions *P* (*ϕ*) ∈ [− *π*, +*π*] at **(a)** the first echo for *f*_*NG*_ and *f*_*G*_, and **(b)** at the final echo for *f*_*NG,NG*_, *f*_exch_, and *f*_*G,G*_ in simulation data generated from a single repetition using *ρ*_mid_ ≈ 1.397 and *t*_*m*_ = 50 ms. The signal *S* = ⟨cos (*ϕ*)⟩ (i.e., real part) is shown as text. The non-exchanging, restricted signal *f*_*NG,NG*_ is well-described by a Gaussian (green, dash-dot line) such that the GPA holds, as expected for this value of *ρ* ≳ 1. Quantitatively, an Anderson-Darling test yields a *p*-value ≈ 0.33. The Gaussian signal *f*_*G,G*_ is fully dephased. The exchanged signal *f*_exch_ is mostly, though not entirely dephased.

Although the GPA holds in the simulation data, the stationarity requirement actually does not hold. The simulation is initialized with a uniform distribution of walkers, which leads to greater exchange out of the sphere(s) compared to inwards (by a factor of about ≈ 10 ×, data not shown). This is simply because the probability of sphere wall collision is much higher for walkers within the sphere than outside. As such, we cannot apply these time-dependent signal models to the simulation DEXR data, though we stress that this does not affect the validity of the previous analyses (Fig. 5) because those were based simply on a quasi-biexponential model of the signal with *f*_*G*_, *f*_*NG*_ and exchange between these pools. Adapting simulations for this time-dependent analysis remains a topic for future work. Nonetheless, we can use the simulation data to form initial intuition about the various transport quantities.

### 4.3. Time-dependent diffusion from simulation

Let us first look at the behavior of the MSD, VACF, *D t*, and *D*_inst_(*t*) from simulation as a representative system with restriction and exchange. In Fig. 10a, we plot the MSD obtained from simulation for times up to *t* = 52.36 ms (*t*_*m*_ = 50, *τ* = 0.59 ms) along with expressions that describe the short- and long-time scaling behaviors. At short times *t* ≪ 0.1 ms, the MSD follows the expected free behavior of 2*D*_0_*t*, but very quickly diverges as walkers interact with walls, taking a concave-down shape. At the tail-end of the simulated range of times, the MSD is better described by an exponential function. In Fig. 10b, we plot the same data and expressions on a log-log plot. To make possible the analysis of the derivative and curvature of the MSD (i.e., to make the MSD smooth and twice-differentiable in order to reveal the VACF), we fit a piecewise, cubic Hermite polynomial [99] to the MSD in this log-log domain (see caption). In Fig. 10c, we plot *D*_inst_ (*t*), estimated by taking a backwards, first-order finite difference of the fitted MSD with time spacing Δ*t* = 5 × 10^− 4^ ms. We also plot *D*(*t*) = ⟨*r*^2^ (*t*)⟩/2*t*, obtained from the raw MSD. Both diffusivity-type quantities are seen to decay monotonically from *D*_0_ = 2.15 *μ*m^2^/ms, which is consistent with the sub-diffusive behavior observed in the MSD.

**Figure 10:**
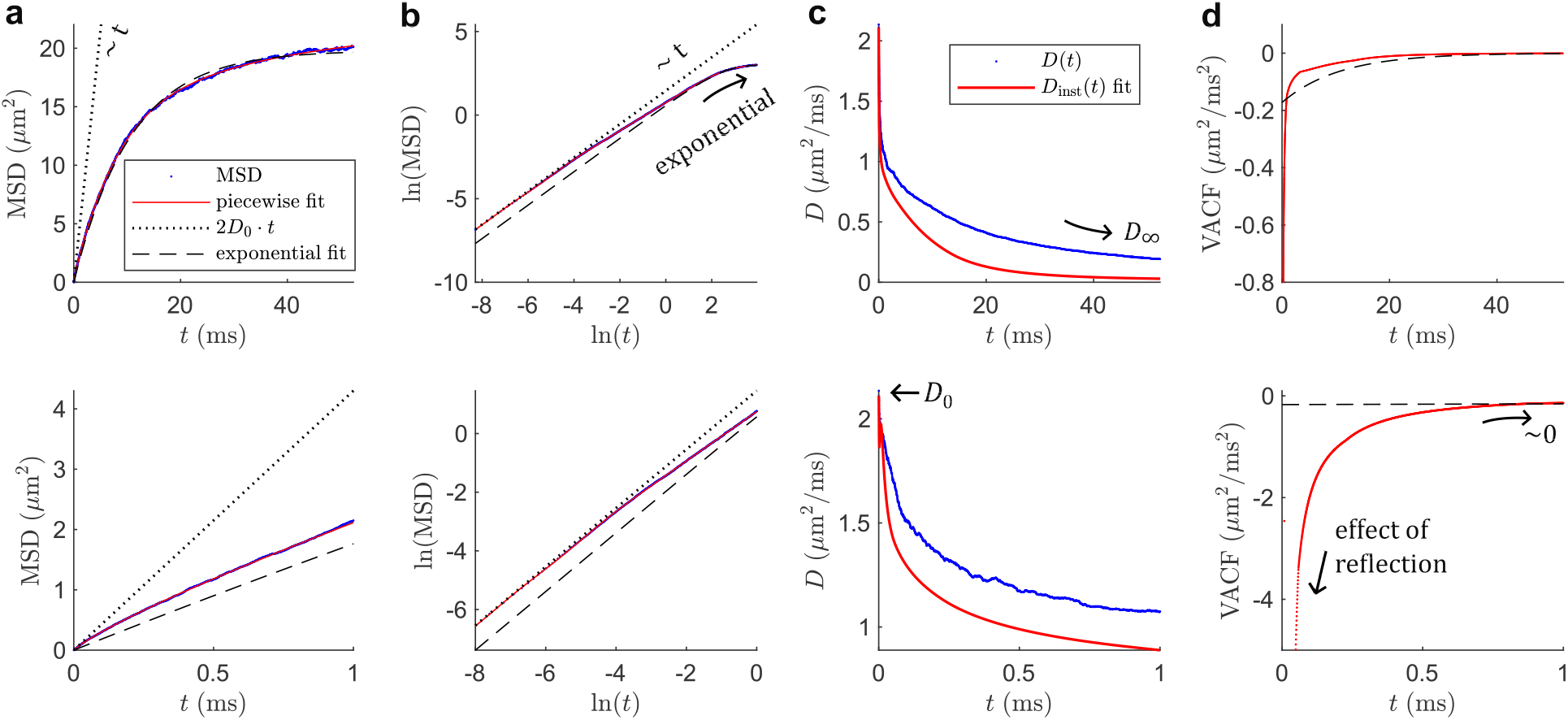
MSD and other transport quantities from simulation. **(a)** The MSD along the gradient direction from one simulation up to *t* ≈ 52.36 ms (blue dots), corresponding to *t*_*m*_ = 50, *τ* = 0.59 ms. Also shown is a linear relationship with *t* (dotted) and an exponential fit (dashed) that describe the MSD at shorter and longer times, respectively. To analyze the first and second derivatives of the MSD, a piecewise, cubic Hermite polynomial was fit in the log-log domain (solid line), splitting the domain into 10 log-linearly spaced segements. For all panels, the bottom plot shows the short-time behavior *t* ≤ 1 ms. Note the immediate deviation from 2*D*_0_*t*. **(b)** Same data and relationships on a log-log plot. The approach towards exponential behavior is clear. **(c)** Diffusivity quantities derived from the MSD. The time-dependent diffusivity *D* (*t*) is the raw MSD divided by 2*t*. The instantaneous diffusivity *D*_inst_ (*t*) is estimated using a backward, first-order finite difference of the piecewise fit to the MSD with spacing Δ*t* = 5 × 10^− 4^ ms, or twice the simulation time-step. Both quantities decay monotonically from *D*_0_. *D* (*t*) approaches a Gaussian limit described by an unknown *D*_∞_ where as *D*_inst_ (*t*) approaches 0, consistent with the bounding box in the simulation. **(d)** The VACF is estimated as half the curvature in the piecewise fit to the MSD obtained using a central, second-order finite difference with the same spacing Δ*t* = 5 × 10^− 4^ ms. The VACF exhibits a sharp, initial decrease due to reflection before asymptotically approaching 0 as *t* → ∞ and the system loses its “memory” of the first interaction(s) with barriers via exchange.

Finally, we plot the VACF in Fig. 10d, estimated by taking a central, second-order finite difference of the fitted MSD using the same Δ*t* = 5 × 10^− 4^ ms spacing. In principle, the VACF should be 0 at *t* = 0 as the MSD is linear and there is no correlation between walker steps. Here, the VACF has decreased rapidly on a timescale that cannot be observed (i.e., time to first interaction with a barrier). This initial decrease in the VACF can be interpreted as the effect of reflection: a walker’s velocity will be negatively correlated with its initial trajectory towards a barrier. Following this decrease, the VACF rises asymptotically towards 0, consistent with “memory” loss of the system, i.e., the de-correlation of walker velocities over time due to exchange. This behavior can be seen in Fig. 10d.

The shape of the VACF here informs the expected behavior in experimental estimates of the VACF using Eq. (45). Because we can only probe *t* > 2*τ*, the short-time decrease in the VACF is not visible, and only the intermediate-to long-time regime over which the VACF approaches 0 can be observed. Therefore, the rate of decay in ln (*S*_mid_/*S*_end_) is related to the rate of growth in the VACF. This relationship is intuitive: if exchange is slow, then walkers that remain confined will exhibit persistent negative autocorrelation(s), slowing the growth of the VACF; if exchange is fast, then velocities will rapidly de-correlate as walkers enter the freer space, increasing the VACF towards 0. Restriction size and shape will also influence the VACF. Smaller restrictions, for example, would result in greater initial decrease of the VACF (i.e., more reflections per unit time, all else being equal). Exchange can thus be thought of as giving rise to or arising from the asymptotic tail of the VACF, with exchange leading to faster recovery. This tail is what is measured using the DEXR method when viewed from the perspective of time-dependent diffusion.

### 4.4. Measuring the VACF

While Eq. (45) is attractive in its simplicity, applying it to actual measurements of *S*_mid_ and *S*_end_ is not straight-forward. Again, the imperfect cancellation of the first lobe(s) in Fig. 8 leads to the constant *C*_0_ in Eq. (44) which we argued can be estimated from *β*_3_ as given in Eq. (18). This may be practically difficult, however, if the data deviates significantly from a first-order exchange model. For instance, we noted some multiexponential character in the spinal cord data in Fig. 6b, indicating that Eq. (18) may not be a sufficient model to describe the signal. There are also errors introduced by finite differencing (on the order of the spacing, 4*τ*/3), as well as blurring of variation in the VACF due to the broadness of the peaks seen in Fig. 8 (width of 2*τ*). This blurring is particularly problematic in the short-time regime (∼ 1 ms) where the VACF changes rapidly (Fig. 10d). Experimental estimates in this regime may flatten the true variation. Furthermore, the exchange weighting in *S*_end_ in acquired data means that 𝒞_end_ (*t*) will also have smaller lobes about *t* = 2*τ*+ *t*_*m*_ such that an additional scaling factor < 1 is necessary to yield the correct proportionality with the VACF. We can nonetheless make a similar argument to that made for measuring *τ*_*k*_: regardless of these other effects, the overall *scaling* behavior w.r.t. *t*_*m*_ should approximate the scaling of the VACF.

What should this behavior be? According to Novikov *et al*. [54], the “structural disorder” of a system, which can be thought of as the distribution of domains or barriers (i.e., their spatial Fourier transform), determines the behavior. Different power-law scaling exponents of ∼ *t*^− *ϑ*^ were proposed in the decay in *D*_inst_ (*t*) as *t* → ∞, corresponding to different “structural universality classes”. From Eq. (36), this corresponds to recovery in the VACF with ∼ *t*^− *ϑ* − 1^. For fully periodic domains, *ϑ* → ∞, and the decay is exponential and thus faster than any power law because walkers do not need to explore the whole domain to reach the limiting Gaussian behavior with VACF = 0. Note that this is consistent with Fig. 10d, where the long-time behavior in the simulation VACF (which has periodic domains) is well-approximated by an exponential. For fully *un*correlated domains, *ϑ* = *d*/2, where *d* is the dimensionality. Other cases with analytical results are random membranes with *ϑ* = 1/2 and random rods with *ϑ* = 1.

In Fig. 11, we plot the apparent VACF estimated by applying Eq. (45) to DEXR data from viable spinal cord. A fit to Eq. (18) was first performed to estimate *β*_3_. To look at the scaling behavior with time, we show a (natural) log-log plot in the insets, taking the absolute value to yield a decay in real values. Also marked is the value of *τ*_*k*_ = 11.7 ms (see Table 1. We plot only up to *t*_*m*_ = 80 ms, as the data reaches a noise floor indistinguishable from 0 at higher *t*_*m*_. The general behavior is as expected, with a monotone approach towards 0 (compare to Fig. 10d). In the log-log inset, it is clear that the approach to 0 sharply accelerates as *t* > *τ*_*k*_, indicative of exchange being the controlling factor in the VACF tail. Thus, the apparent VACF curve reproduces expected trends. As an initial sanity check, numerically integrating the apparent VACF yields ≈ − 0.8 *μ*m^2^/ms, corresponding to a decrease in *D*_inst_ (*t*) from ≈ 2.15 → 1.35 *μ*m^2^/ms over the observed time frame. This is roughly the expected magnitude of decrease (i.e., decreases less than *D*_0_), given that some further negative portion of the VACF is not visible at short times.

**Figure 11:**
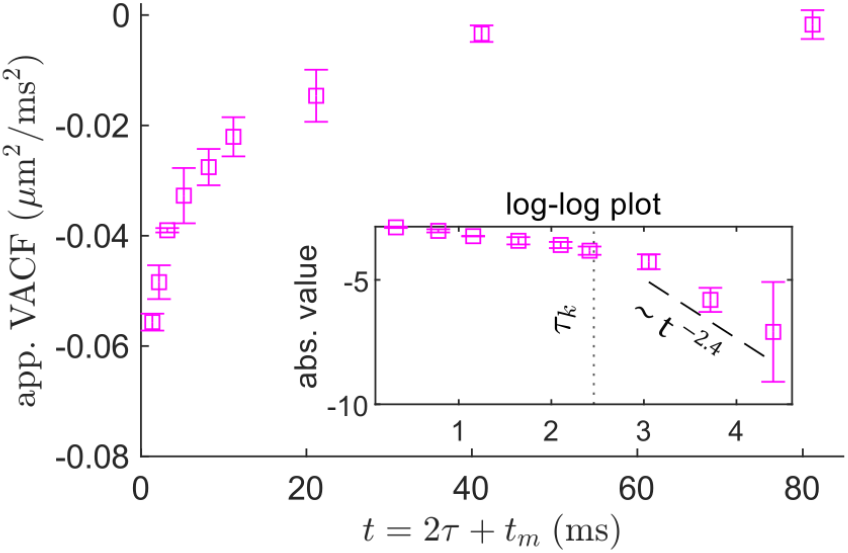
Apparent VACF from DEXR data viable spinal cord, calculated using Eqs. (45) and (18) to obtain *β*_3_. Error bars = mean ± SD from repetitions with different seeds. For each repetition, *β*_3_ was first estimated by fitting to Eq. (18) and the VACF was then calculated as 3/(2*τb*_*s*_)[ln (*β*_3_) − ln (*S*_mid_/*S*_end_)]as in Eq. (45). The leading factor is calculated as 3/(2*τb*_*s*_) ≈ 0.55 *μ*m^2^/ms^2^ using *b*_*s*_ ≈ 4.59 ms/*μ*m^2^ and *τ* = 0.59 ms. Insets show a (natural) log-log plot obtained by first taking the absolute value of the VACF. The exchange time, *τ*_*k*_ = 11.7 ms is marked with a dotted line. A subset of the data is shown, omitting the longest mixing times at *t*_*m*_ = [160, 300] ms that are effectively 0. In the log-log inset, a linear fit to the points over *t*_*m*_ = [20, 40, 80] (dashed line) indicates a power-law tail with ∼ *t* ^− 2.4^, or *ϑ* = 1.4, although this fit is highly sensitive to *β*_3_.

What of the scaling? In the log-log inset of Fig. 11, we perform a linear fit in this domain over several mixing times for which *t* > *τ*_*k*_ at *t*_*m*_ = [20, 40, 80] ms (dashed line), yielding a scaling with ∼ *t*^− 2.4^ or *ϑ* ≈ 1.4. This lies between the exponents for uncorrelated domains in 3-D (*ϑ* = 3/2) and random rods (*ϑ* = 1), which may indeed be consistent with the makeup of GM (i.e., soma and neurites). This differs from the *ϑ* ≈ 1/2 estimated by Novikov *et al*. in GM [54, 19], which they argued is consistent with uncorrelated domains in 1-D. A potential biological substrate of such behavior is beads or varicosities [100] along effectively 1-D neurites. Here, the short dephasing length *𝓁*_*g*_ may restore the intrinsic 3-D nature of neurites (i.e., we may be sensitive to decay due to diffusion in the intra-neurite space), resulting in *ϑ* = 3/2. In the study by Novikov *et al*. [54], however, the data was found to follow a single power law across *ω* ≈ 0 − 500 Hz or from *t* ≈ 2 ms to long times. We do not see this behavior here, where there is a clear transition due to exchange.

As a precaution, we stress that estimates of *ϑ* are highly sensitive to the estimate of *β*_3_ (data not shown) — i.e., a smaller *β*_3_ would yield a potentially much slower decay in the log-log domain — and this estimate relies on a perhaps flawed assumption of monoexponential behavior in the data, mentioned above. This issue is ameliorated by going to very long *t*_*m*_ ≫ *τ*_*k*_ up to 300 ms ≈ 28*τ*_*k*_ in the spinal cord data, and it may be argued that the estimate of *β*_3_ is robust to the presence of multiexponential behavior unless such behavior manifests over very long times. Furthermore, while *ϑ* is a straightforward observable that can be compared to the literature, we emphasize that the DEXR method yields the apparent VACF outright over a wide range of times, and different analyses or quantitative metrics may also be insightful. As an example, the data suggest that piece-wise fitting of power laws with a transition at *t* = *τ*_*k*_ may fit the VACF well (see again the inset in Fig. 11).

### 4.5. Reconciling the two interpretations

Taking a step back, this alternative interpretation in terms of the VACF sheds light on the behavior of DEXR data and whether they can be described by a first-order exchange model. When modelling exchange, or when using compartment-based signal models in general, it is tempting to argue that deviations can be explained by further compartments and parameters. The VACF provides a more model-agnostic view. The multiexponential character in the spinal cord data seen in Fig. 6b could potentially be described as a result of complicated behavior in the VACF, which could also involve multi-site exchange as hypothesized by Cai *et al*. [90]. By bridging these sub-fields of diffusion MR under one method, we can assess how phenomena such as exchange are related to fundamental transport quantities such as the MSD. Instead of assuming some compartmentalization, we can instead begin from models of the MSD and make forward predictions of the VACF tail and DEXR data.

For instance, we can compare to the literature on anomalous diffusion modelling (see ref. [101] for brief review) wherein power laws are used to describe the MSD. Because the VACF is the curvature in the MSD, see again Eq. (36), these exponents in the MSD translate directly to power law scaling in the VACF by an exponent subtracted by 2. Other models of the MSD include the Ornstein-Uhlenbeck model [102], which predicts exponential recovery in the MSD and thus corresponding exponential behavior in the VACF [55]. Many such analyses that begin with an analytical form of the MSD are possible, and we leave this as a topic for future work.

## 5. Discussion and conclusions

### 5.1. Summary of findings

This work provides theoretical underpinnings and guidelines for the design, optimization, and data interpretation of a two-point SG-DEXSY sub-sampling scheme, which we call the DEXR method. Based on taking the ratio of equally diffusion-weighted, but oppositely exchange-weighted points — *S*_mid_ and *S*_end_ — the method was shown to produce robust estimates of *τ*_*k*_ and restriction parameters in simulation data. The method was subsequently applied in viable, *ex vivo* spinal cord of neonatal mice using a high gradient system, yielding *τ*_*k*_ ≈ 11 ms, *f*_*I*_ ≈ 0.71, and *R*_eff_ ≈ 1.1 *μ*m from just 6 total data points. Our findings highlight the specificity as well as efficiency of the method for probing microstructural features. Importantly, the method decouples the measurement of exchange from restriction and overcomes this degeneracy. Taking a different view, we show that DEXR data can be interpreted to yield an apparent VACF. To our knowledge, this is the only method capable of yielding point-wise sampling in the time-domain without the use of oscillating gradients. The DEXR method enables the study of the VACF across a wide range of times (*t* ∼ 2 − 500 ms) while using the same experimental paradigm. Preliminarily, we find long-time scaling behavior (*t* ∼ 20 − 80 ms) in viable spinal cord that is roughly consistent with short-range, 3-D disorder (*ϑ* ≈ 3/2).

### 5.2. Limitations and assumptions

We were careful throughout to state the assumptions required for each analysis. For instance, we found that we could not yield an apparent VACF from simulation data due to non-stationarity. The assumptions required to estimate restriction parameters are particularly nuanced, with various corrections (*σ, ς, η*) needed to yield quantitative *f*_*I*_ and ⟨*c*_*I*_⟩ values. The downstream estimation of *R*_eff_ and *κ*_eff_ requires an additional assumption of spherical compartments (without localized signal), which may not be accurate. Note that changing to other effective geometries (cylindrical, parallel plates) merely involves changing the constant prefactor in Eq. (9). With regards to exchange, we argued that the disparate *τ*_*k*_ estimates in the literature [86] can perhaps be explained by differences in *𝓁*_*g*_ (see Sec. 3.1) and will not remark further. Some of our assumptions, however, merit reexamination. Most notably, we ignored *T*_2_ relaxation and *T*_2_-*T*_2_ exchange effects [88, 103] on the basis of short diffusion encodings, *τ* < 1 ms. For systems with smaller gradient amplitude, longer encoding times become necessary to reach optimal exchange weighting (see Figs. 3 and 4) and these effects may become significant. For PG systems, these effects can perhaps be normalized by using fixed diffusion times (not possible on our SG system), but this changes the relevant theory as *𝓁*_*g*_ is now variable, which we discuss in the following subsection. Other effects that were neglected include surface relaxation and magnetization transfer [104], though we suspect that these effects will manifest in the effective diffusion-weighted relaxation rates such that they do not need to be explicitly included in our signal model(s).

An issue that is more difficult to address is the possible breakdown of detailed balance. Investigations of relaxation exchange [105, 106] find that the exchange map in multi-site (> 2) exchange can be asymmetric, indicative of a circular exchange pathway. Such exchange pathways would complicate our view of the Gaussian and non-Gaussian pools being static over time and *f*_exch_ may exhibit unexpected decay behavior. While a breakdown of detailed balance has not yet been demonstrated in diffusion MR data (to our knowledge) this cannot be excluded as a possibility, and may lead to bias in our estimates of *τ*_*k*_. A similar issue is the breakdown of first-order exchange. Such a breakdown was recently discussed by Ordinola *et al*. [89] in the context of a discrete diffusion spectrum. This was also observed in Cai *et al*. [90], and both works report multiexponential behavior in the exchange-weighted signal measured via DEXSY. We add our own results regarding the VACF as a possible explanation for this behavior (see Fig. 6b), and reiterate that a first-order exchange model is not necessarily compatible with what is seen in the VACF, though empirical agreement is observed here (see again Figs. 5, 6b, 11). This also calls into question whether the wider body of literature (e.g., NEXI [29]) based on the first-order Kärger model [9, 10] may be affected by the breakdown of first-order exchange.

### 5.3. Application to pulsed gradients

The DEXR method and framework can readily be applied to PG experiments if the separation between gradient lobes Δ − *δ* is small compared to the gradient duration *δ*, and furthermore the diffusion weighting is varied by changing the timings, rather than the gradient amplitude *g*. Such an experiment resembles the SG case, and the same principles can be applied. Typical PG experiments, however, are not performed in this way and instead fix the timings *δ*, Δ while varying *g*. In this PG case, we cannot easily condense the experimental parameters by defining *ρ* := *𝓁*_*d*_/*𝓁*_*g*_ as we did for the SG case. Consider for instance that the motionally-averaged signal behavior in Eq. (4) would become

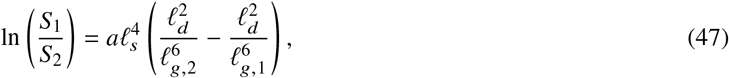

where *S*_1_ and *S*_2_ are two acquisitions with different *𝓁*_*g*,1_ and *𝓁*_*g*,2_, and 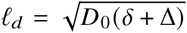. The expression cannot be condensed due to the different powers in *𝓁*_*d*_ and *𝓁*_*g*_. The sensitivity to exchange would also differ between acquisitions, following our argument in Sec. 3.1. It may thus be more difficult to extract restriction and exchange parameters as at least one additional parameter is needed in the signal model. On the other hand, the normalization of *T*_2_ relaxation effects becomes more straightforward, as *𝓁*_*d*_ is not varied. For relatively moderate *b*-values ∼ 4 ms/*μ*m^2^ such as those used here, the necessary difference in *𝓁*_*g*,1_ and *𝓁*_*g*,2_ to yield the same total *b*_*s*_ (i.e., to acquire *S*_mid_ and *S*_end_) may be small, and the signal models presented in this paper may be sufficient to a first approximation.

### 5.4. Comparison to related work

Our method is innovative in its analysis but its methodology is similar to other approaches based on DEXSY and/or on the ratio of acquisitions with different diffusion contrasts. The most relevant point of comparison is to FEXSY [41], which, similar to our method, is based on a sub-sampling of DEXSY data and does not employ a numerical inverse Laplace transform. Practically, our method reduces to FEXSY when the filter value of *b*_1_ is set to be equal to *b*_2_ and the baseline ADC is then measured using *b*_*s*_ = *b*_1_ + *b*_2_. Indeed, a similar approach to fitting the exchange time was presented by Scher *et al*. [107], also using constant gradients. We stress, however, that the FEXSY model is fundamentally different in that it looks at an ADC recovery, which does not account for non-Gaussian diffusion. The downstream analyses to yield *f*_*I*_ and *R*_eff_ are unique to this work, as is the estimation of the VACF. FEXSY also does not account for a diffusion-weighted *T*_1_ in its original conception.

Parallels can also be drawn to the “temporal diffusion ratio” (TDR) described recently by Warner *et al*. [108]. In TDR, the ratio of two acquisitions with varying diffusion times but fixed total *b*-value is assessed to yield microstructural contrast, similar to *S*_mid_ and *S*_end_ acquired here. Like FEXSY, however, TDR produces an empirical contrast between these acquisitions, without attempting to extract quantitative microstructural parameters. Once again, it is the modelling and analysis that separate the DEXR method and make it uniquely information-rich. A further advantage of DEXR is its ability to vary the time-domain weighting via *t*_*m*_, which gives a wide range of time sensitivity compared to TDR and even TDS based on oscillating gradients. We also point out that while stimulated echoes have been used as a means to probe long-time diffusion (e.g., by Fieremans *et al*. [109]), the time-domain weighting of a typical diffusion-weighted stimulated echo as given by 𝒞(*t*) in Eq. (38) would be very broad, spanning all observed times. As such, the ability to resolve long-time behavior using this approach is limited. DEXR has the advantage of truly isolating the variation at long diffusion times.

### 5.5. Concluding remarks

While challenges remain, particularly in adapting the DEXR method to PG experiments and in validating VACF measurements, we demonstrate in this work that the method can yield quantitative exchange, restriction, and time-dependence information from sparse diffusion MR data. Compared to other approaches, the method is highly specific and efficient. We have provided herein a thorough description, validation (via simulation), and proof-of-concept (mouse spinal cord, with the PM-10) for the DEXR method and pave the way for future applications. The method may be especially useful in hitherto difficult to characterize samples that have overlapping exchange and restriction effects such as GM.

## Data Availability Statement

Spinal cord data and NMR sequences, as well as Julia code to run the Monte Carlo simulations and MATLAB fitting routines are available upon reasonable request.

## CRediT Author Contributions

**Teddy X. Cai:** Conceptualization, Data Curation, Formal Analysis, Methodology, Software, Validation, Visualization, Writing – Original Draft, Writing – Review & Editing. **Nathan H. Williamson:** Conceptualization, Data Curation, Methodology, Investigation, Resources, Writing - Original Draft, Writing – Review & Editing. **Rea Ravin:** Methodology, Investigation, Resources, Writing - Review & Editing. **Peter J. Basser:** Supervision, Writing - Review & Editing.

## Acknowledgements

The authors would like to thank Drs. Michael O’Donovan and Melanie Falgairolle for their help with *ex vivo* tissue preparation protocols.

## References

[1] D. E. Woessner, Nmr spin-echo self-diffusion measurements on fluids undergoing restricted diffusion, The Journal of Chemical Physics 67 (6) (1963) 1365–1367. doi:10.1021/j100800a509.

[2] E. O. Stejskal, Use of spin echoes in a pulsed magnetic-field gradient to study anisotropic, restricted diffusion and flow, The Journal of Chemical Physics 43 (10) (1965) 3597–3603. doi:10.1063/1.1696526.

[3] B. Robertson, Spin-echo decay of spins diffusing in a bounded region, Physical Review 151 (1966) 273–277. doi:10.1103/PhysRev.151.273.

[4] J. E. Tanner, E. O. Stejskal, Restricted self-diffusion of protons in colloidal systems by the pulsed-gradient, spin-echo method, The Journal of Chemical Physics 49 (4) (1968) 1768–1777. doi:10.1063/1.1670306.

[5] C. H. Neuman, Spin echo of spins diffusing in a bounded medium, The Journal of Chemical Physics 60 (11) (1974) 4508–4511. doi:10.1063/1.1680931.

[6] P. P. Mitra, P. N. Sen, L. M. Schwartz, P. Le Doussal, Diffusion propagator as a probe of the structure of porous media, Physical Review Letters 68 (1992) 3555–3558. doi:10.1103/PhysRevLett.68.3555.

[7] L. L. Latour, P. P. Mitra, R. L. Kleinberg, C. H. Sotak, Time-dependent diffusion coefficient of fluids in porous media as a probe of surface-to-volume ratio, Journal of Magnetic Resonance 101 (3) (1993) 342–346. doi:10.1006/jmra.1993.1056.

[8] J. E. Tanner, Transient diffusion in a system partitioned by permeable barriers. application to nmr measurements with a pulsed field gradient, The Journal of Chemical Physics 69 (4) (1978) 1748–1754. doi:10.1063/1.436751.

[9] J. Kärger, Zur bestimmung der diffusion in einem zweibereichsystem mit hilfe von gepulsten feldgradienten, Annalen der Physik 479 (1–2) (1969) 1–4. doi:10.1002/andp.19694790102.

[10] J. Kärger, Nmr self-diffusion studies in heterogeneous systems, Adv. Colloid Interface Sci. 23 (1985) 129–148. doi:10.1016/0001-8686(85)80018-X.

[11] L. L. Latour, K. Svoboda, P. P. Mitra, C. H. Sotak, Time-dependent diffusion of water in a biological model system, Proceedings of the National Academy of Sciences 91 (4) (1994) 1229–1233. doi:10.1073/pnas.91.4.1229.

[12] P. N. Sen, Time-dependent diffusion coefficient as a probe of the permeability of the pore wall, The Journal of Chemical Physics 119 (18) (2003) 9871–9876. doi:10.1063/1.1611477.

[13] D. S. Novikov, E. Fieremans, J. H. Jensen, J. A. Helpern, Random walk with barriers, Nature Physics 7 (6) (2011) 508–514. doi:10.1038/nphys1936.

[14] E. O. Stejskal, J. E. Tanner, Spin diffusion measurements: Spin echoes in the presence of a time-dependent field gradient, The Journal of Chemical Physics 42 (1) (1965) 288–292. doi:10.1063/1.1695690.

[15] I. O. Jelescu, J. Veraart, E. Fieremans, D. S. Novikov, Degeneracy in model parameter estimation for multi-compartmental diffusion in neuronal tissue, NMR in Biomedicine 29 (1) (2015) 33–47. doi:10.1002/nbm.3450.

[16] I. O. Jelescu, M. Palombo, F. Bagnato, K. G. Schilling, Challenges for biophysical modeling of microstructure, Journal of Neuroscience Methods 344 (2020) 108861. doi:10.1016/j.jneumeth.2020.108861.

[17] D. S. Novikov, J. Veraart, I. O. Jelescu, E. Fieremans, Rotationally-invariant mapping of scalar and orientational metrics of neuronal microstructure with diffusion mri, NeuroImage 174 (2018) 518—-538. doi:10.1016/j.neuroimage.2018.03.006.

[18] D. S. Novikov, V. G. Kiselev, S. N. Jespersen, On modeling, Magnetic Resonance in Medicine 79 (6) (2018) 3172–3193. doi:10.1002/mrm.27101.

[19] D. S. Novikov, E. Fieremans, S. N. Jespersen, V. G. Kiselev, Quantifying brain microstructure with diffusion mri: Theory and parameter estimation, NMR in Biomedicine 32 (4) (2019) e3998. doi:10.1002/nbm.3998.

[20] S. Coelho, J. M. Pozo, S. N. Jespersen, D. K. Jones, A. F. Frangi, Resolving degeneracy in diffusion MRI biophysical model parameter estimation using double diffusion encoding, Magnetic Resonance in Medicine 82 (1) (2019) 395–410. doi:10.1002/mrm.27714.

[21] A. Chakwizira, C.-F. Westin, J. Brabec, S. Lasič, L. Knutsson, F. Szczepankiewicz, M. Nilsson, Diffusion mri with pulsed and free gradient waveforms: Effects of restricted diffusion and exchange, NMR in Biomedicine 36 (1) (2023) e4827. doi:10.1002/nbm.4827.

[22] M. C. Papadopoulos, A. S. Verkman, Aquaporin water channels in the nervous system, Nature Reviews Neuroscience 14 (4) (2013) 265–277. doi:10.1038/nrn3468.

[23] N. H. Williamson, R. Ravin, D. Benjamini, H. Merkle, M. Falgairolle, M. J. O’Donovan, D. Blivis, D. Ide, T. X. Cai, N. S. Ghorashi, R. Bai, P. J. Basser, Magnetic resonance measurements of cellular and sub-cellular membrane structures in live and fixed neural tissue, eLife 8 (2019) e51101. doi:10.7554/eLife.51101.

[24] H. H. Lee, J. L. Olesen, Q. Tian, G. R. Llorden, S. N. Jespersen, S. Y. Huang, Revealing diffusion time-dependence and exchange effect in the in vivo human brain gray matter by using high gradient diffusion mri, in: Proceedings of the Annual Meeting of the International Society of Magnetic Resonance in Medicine, Vol. 30, 2022, p. 0254.

[25] J. L. Olesen, L. Østergaard, N. Shemesh, S. N. Jespersen, Diffusion time dependence, power-law scaling, and exchange in gray matter, NeuroImage 251 (2022) 118976. doi:10.1016/j.neuroimage.2022.118976.

[26] G. J. Stanisz, G. A. Wright, R. M. Henkelman, A. Szafer, An analytical model of restricted diffusion in bovine optic nerve, Magnetic Resonance in Medicine 37 (1) (1997) 103—-111. doi:10.1002/mrm.1910370115.

[27] J. Pfeuffer, U. Flögel, W. Dreher, D. Leibfritz, Restricted diffusion and exchange of intracellular water: theoretical modelling and diffusion time dependence of 1h nmr measurements on perfused glial cells, NMR in Biomedicine 11 (1) (1998) 19––31. doi:10.1002/(sici)1099-1492(199802)11:1¡19::aid-nbm499¿3.0.co;2-o.

[28] K. J. Carlton, M. R. Halse, J. H. Strange, Diffusion-weighted imaging of bacteria colonies in the STRAFI plane, Journal of Magnetic Resonance 143 (1) (2000) 24–29. doi:10.1006/jmre.1999.1959.

[29] I. O. Jelescu, A. de Skowronski, F. Geffroy, M. Palombo, D. S. Novikov, Neurite exchange imaging (nexi): A minimal model of diffusion in gray matter with inter-compartment water exchange, NeuroImage 256 (2022) 119277. doi:10.1016/j.neuroimage.2022.119277.

[30] M. Palombo, A. Ianus, M. Guerreri, D. Nunes, D. C. Alexander, N. Shemesh, H. Zhang, Sandi: A compartment-based model for non-invasive apparent soma and neurite imaging by diffusion mri, NeuroImage 215 (2020) 116835. doi:10.1016/j.neuroimage.2020.116835.

[31] Q. Uhl, T. Pavan, M. Molendowska, D. K. Jones, M. Palombo, I. O. Jelescu, Quantifying human gray matter microstructure using neurite exchange imaging (nexi) and 300 mt/m gradients, Imaging Neuroscience 2 (2024) 1–19. doi:10.1162/imag.a.00104.

[32] R. N. Henriques, M. Palombo, S. N. Jespersen, N. Shemesh, H. Lundell, A. Ianuş, Double diffusion encoding and applications for biomedical imaging, Journal of Neuroscience Methods 348 (2021) 108989. doi:10.1016/j.jneumeth.2020.108989.

[33] P. T. Callaghan, I. Furó, Diffusion-diffusion correlation and exchange as a signature for local order and dynamics, The Journal of Chemical Physics 120 (8) (2004) 4032–4038. doi:10.1063/1.1642604.

[34] J. O. Breen-Norris, B. Siow, C. Walsh, B. Hipwell, I. Hill, T. Roberts, M. G. Hall, M. F. Lythgoe, A. Ianus, D. C. Alexander, S. Walker-Samuel, Measuring diffusion exchange across the cell membrane with dexsy (diffusion exchange spectroscopy), Magnetic Resonance in Medicine 84 (3) (2020) 1543–1551. doi:10.1002/mrm.28207.

[35] D. Benjamini, M. E. Komlosh, P. J. Basser, Imaging local diffusive dynamics using diffusion exchange spectroscopy mri, Physical Review Letters 118 (2017) 158003. doi:10.1103/PhysRevLett.118.158003.

[36] O. Mankinen, V. V. Zhivonitko, A. Selent, S. Mailhiot, S. Komulainen, N. L. Prisle, S. Ahola, V.-V. Telkki, Ultrafast diffusion exchange nuclear magnetic resonance, Nature Communications 11 (1) (2020) 3251. doi:10.1038/s41467-020-17079-7.

[37] Y. Qiao, P. Galvosas, T. Adalsteinsson, M. Schönhoff, P. T. Callaghan, Diffusion exchange nmr spectroscopic study of dextran exchange through polyelectrolyte multilayer capsules, The Journal of Chemical Physics 122 (21) (2005). doi:10.1063/1.1924707.

[38] R. Bai, A. Cloninger, W. Czaja, P. J. Basser, Efficient 2d mri relaxometry using compressed sensing, Journal of Magnetic Resonance 255 (2015) 88–99. doi:10.1016/j.jmr.2015.04.002.

[39] R. Bai, D. Benjamini, J. Cheng, P. J. Basser, Fast, accurate 2D-MR relaxation exchange spectroscopy (REXSY): Beyond compressed sensing, The Journal of Chemical Physics 145 (15) (2016) 154202. doi:10.1063/1.4964144.

[40] D. Benjamini, P. J. Basser, Use of marginal distributions constrained optimization (MADCO) for accelerated 2d MRI relaxometry and diffusometry, Journal of Magnetic Resonance 271 (2016) 40–45. doi:10.1016/j.jmr.2016.08.004.

[41] I. Åslund, A. Nowacka, M. Nilsson, D. Topgaard, Filter-exchange PGSE NMR determination of cell membrane permeability, Journal of Magnetic Resonance 200 (2) (2009) 291–295.

[42] T. X. Cai, D. Benjamini, M. E. Komlosh, P. J. Basser, N. H. Williamson, Rapid detection of the presence of diffusion exchange, Journal of Magnetic Resonance 297 (2018) 17–22. doi:10.1016/j.jmr.2018.10.004.

[43] N. H. Williamson, R. R. T. X. Cai, D. Benjamini, M. Falgairolle, M. J. O’Donovan, P. J. Basser, Real-time measurement of diffusion exchange rate in biological tissue, Journal of Magnetic Resonance 317 (2020) 106782. doi:10.1016/j.jmr.2020.106782.

[44] J. E. Tanner, Self diffusion of water in frog muscle, Biophys. J. 28 (1) (1979) 107–116. doi:10.1016/s0006-3495(79)85162-0.

[45] J. StepiŞnik, Analysis of nmr self-diffusion measurements by a density matrix calculation, Physica B+C 104 (3) (1981) 350–364. doi:10.1016/0378-4363(81)90182-0.

[46] J. StepiŞnik, Time-dependent self-diffusion by nmr spin-echo, Physica B Condensed Matter 183 (4) (1993) 343–350. doi:10.1016/0921-4526(93)90124-O.

[47] P. T. Callaghan, J. StepiŞnik, Frequency-domain analysis of spin motion using modulated-gradient nmr, Journal of Magnetic Resonance 117 (1) (1995) 118–122. doi:10.1006/jmra.1995.9959.

[48] E. C. Parsons Jr., M. D. Does, J. C. Gore, Temporal diffusion spectroscopy: Theory and implementation in restricted systems using oscillating gradients, Magnetic Resonance in Medicine 55 (1) (2006) 75–84. doi:10.1002/mrm.20732.

[49] J. C. Gore, J. Xu, D. C. Colvin, T. E. Yankeelov, E. C. Parsons, M. D. Does, Characterization of tissue structure at varying length scales using temporal diffusion spectroscopy, NMR in Biomedicine 23 (7) (2010) 745–756. doi:10.1002/nbm.1531.

[50] H. Li, J. C. Gore, J. Xu, Fast and robust measurement of microstructural dimensions using temporal diffusion spectroscopy, Journal of Magnetic Resonance 242 (2014) 4 – 9. doi:10.1016/j.jmr.2014.02.007.

[51] X. Jiang, H. Li, J. Xie, P. Zhao, J. C. Gore, J. Xu, Quantification of cell size using temporal diffusion spectroscopy, Magnetic Resonance in Medicine 75 (3) (2016) 1076–1085. doi:10.1002/mrm.25684.

[52] O. Reynaud, Time-dependent diffusion mri in cancer: Tissue modeling and applications, Frontiers in Physics 5 (2017) 58. doi:10.3389/fphy.2017.00058.

[53] P. P. Mitra, P. N. Sen, L. M. Schwartz, Short-time behavior of the diffusion coefficient as a geometrical probe of porous media, Physical Review B 47 (1993) 8565–8574. doi:10.1103/PhysRevB.47.8565.

[54] D. S. Novikov, J. H. Jensen, J. A. Helpern, E. Fieremans, Revealing mesoscopic structural universality with diffusion, Proceedings of the National Academy of Sciences 111 (14) (2014) 5088–5093. doi:10.1073/pnas.1316944111.

[55] L. Ning, K. Setsompop, C.-F. Westin, Y. Rathi, New insights about time-varying diffusivity and its estimation from diffusion mri, Magnetic Resonance in Medicine 78 (2) (2017) 763–774. doi:10.1002/mrm.26403.

[56] T. X. Cai, N. H. Williamson, V. J. Witherspoon, R. Ravin, P. J. Basser, A single-shot measurement of time-dependent diffusion over sub-millisecond timescales using static field gradient NMR, Journal of Chemical Physics 154 (11) (2021) 111105. doi:10.1063/5.0041354.

[57] E. L. Hahn, Spin echoes, Physical Review 80 (1950) 580–594. doi:10.1103/PhysRev.80.580.

[58] A. M. Henry, J. G. Hohmann, High-resolution gene expression atlases for adult and developing mouse brain and spinal cord, Mammalian Genome 23 (9–10) (2012) 539—-549. doi:10.1007/s00335-012-9406-2.

[59] G. Sengul, R. B. Puchalski, C. Watson, Cytoarchitecture of the spinal cord of the postnatal (p4) mouse, The Anatomical Record 295 (5) (2012) 837—-845. doi:10.1002/ar.22450.

[60] G. Eidmann, R. Savelsberg, P. Blümler, B. Blümich, The nmr mouse, a mobile universal surface explorer, Journal of Magnetic Resonance 122 (1) (1996) 104–109. doi:10.1006/jmra.1996.0185.

[61] B. Blümich, P. Blümler, G. Eidmann, A. Guthausen, R. Haken, U. Schmitz, K. Saito, G. Zimmer, The nmr-mouse: construction, excitation, and applications, Magnetic Resonance Imaging 16 (5) (1998) 479–484. doi:10.1016/S0730-725X(98)00069-1.

[62] S. Utsuzawa, E. Fukushima, Unilateral nmr with a barrel magnet, Journal of Magnetic Resonance 282 (2017) 104–113.

[63] N. H. Williamson, R. Ravin, T. X. Cai, M. Falgairolle, M. J. O’Donovan, P. J. Basser, Water exchange rates measure active transport and homeostasis in neural tissue, PNAS Nexus 2 (3) (2023) pgad056. doi:10.1093/pnasnexus/pgad056.

[64] D. Rata, F. Casanova, J. Perlo, D. Demco, B. Blümich, Self-diffusion measurements by a mobile single-sided nmr sensor with improved magnetic field gradient, Journal of Magnetic Resonance 180 (2) (2006) 229–235.

[65] F. Casanova, J. Perlo, B. Blümich, Single-Sided NMR, Springer Berlin Heidelberg, 2011. doi:10.1007/978-3-642-16307-4.

[66] H. Y. Carr, E. M. Purcell, Effects of diffusion on free precession in nuclear magnetic resonance experiments, Physical Review 94 (1954) 630–638. doi:10.1103/PhysRev.94.630.

[67] S. Meiboom, D. Gill, Modified spin-echo method for measuring nuclear relaxation times, Review of Scientific Instruments 29 (8) (1958) 688–691. doi:10.1063/1.1716296.

[68] M. G. Hall, D. C. Alexander, Convergence and parameter choice for monte-carlo simulations of diffusion mri, IEEE Transactions on Medical Imaging 28 (9) (2009) 1354–1364. doi:10.1109/TMI.2009.2015756.

[69] M. D. Hürlimann, K. G. Helmer, T. M. de Swiet, P. N. Sen, Spin echoes in a constant gradient and in the presence of simple restriction, Journal of Magnetic Resonance, Series A 113 (1995) 260–264. doi:10.1006/jmra.1995.1091.

[70] S. Axelrod, P. N. Sen, Nuclear magnetic resonance spin echoes for restricted diffusion in an inhomogeneous field: Methods and asymptotic regimes, The Journal of Chemical Physics 114 (15) (2001) 6878–6895. doi:10.1063/1.1356010.

[71] M. D. Hürlimann, Diffusion and relaxation effects in general stray field nmr experiments, Journal of Magnetic Resonance 148 (2) (2001) 367–378. doi:10.1006/jmre.2000.2263.

[72] D. S. Grebenkov, Nmr survey of reflected brownian motion, Reviews of Modern Physics 79 (2007) 1077–1137. doi:10.1103/RevModPhys.79.1077.

[73] D. A. Yablonskiy, A. L. Sukstanskii, Theoretical models of the diffusion weighted MR signal, NMR in Biomedicine 23 (7) (2010) 661–681. doi:10.1002/nbm.1520.

[74] H. C. Torrey, Bloch equations with diffusion terms, Physical Review 104 (1956) 563–565. doi:10.1103/PhysRev.104.563.

[75] J. S. Murday, R. M. Cotts, Self-Diffusion Coefficient of Liquid Lithium, The Journal of Chemical Physics 48 (11) (1968) 4938–4945. doi:10.1063/1.1668160.

[76] S. D. Stoller, W. Happer, F. J. Dyson, Transverse spin relaxation in inhomogeneous magnetic fields, Physical Review A 44 (1991) 7459– 7477. doi:10.1103/PhysRevA.44.7459.

[77] T. M. de Swiet, P. N. Sen, Decay of nuclear magnetization by bounded diffusion in a constant field gradient, The Journal of Chemical Physics 100 (8) (1994) 5597–5604. doi:10.1063/1.467127.

[78] N. Moutal, D. S. Grebenkov, The localization regime in a nutshell, Journal of Magnetic Resonance 320 (2020) 106836. doi:10.1016/j.jmr.2020.106836.

[79] D. S. Grebenkov, Diffusion mri/nmr at high gradients: Challenges and perspectives, Microporous and Mesoporous Materials 269 (2018) 79–82. doi:10.1016/j.micromeso.2017.02.002.

[80] N. H. Williamson, V. J. Witherspoon, T. X. Cai, R. Ravin, F. Horkay, P. J. Basser, Low-field, high-gradient nmr shows diffusion contrast consistent with localization or motional averaging of water near surfaces, Magnetic Resonance Letters 3 (2) (2023) 90–107.

[81] T. X. Cai, N. H. Williamson, R. Ravin, P. J. Basser, Disentangling the effects of restriction and exchange with diffusion exchange spectroscopy, Frontiers in Physics 10 (2022). doi:10.3389/fphy.2022.805793.

[82] Y. Assaf, P. J. Basser, Composite hindered and restricted model of diffusion (charmed) mr imaging of the human brain, NeuroImage 27 (1) (2005) 48 – 58. doi:10.1016/j.neuroimage.2005.03.042.

[83] L. M. Burcaw, E. Fieremans, D. S. Novikov, Mesoscopic structure of neuronal tracts from time-dependent diffusion, NeuroImage 114 (2015) 18–37. doi:10.1016/j.neuroimage.2015.03.061.

[84] D. S. Grebenkov, Exploring diffusion across permeable barriers at high gradients. ii. localization regime, Journal of Magnetic Resonance 248 (2014) 164–176. doi:10.1016/j.jmr.2014.08.016.

[85] M. Nilsson, J. Lätt, D. van Westen, S. Brockstedt, S. Lasič, F. Ståhlberg, D. Topgaard, Noninvasive mapping of water diffusional exchange in the human brain using filter-exchange imaging, Magn. Reson. Med. 69 (6) (2013) 1572–1580.

[86] M. Nilsson, D. van Westen, F. Ståhlberg, P. C. Sundgren, J. Lätt, The role of tissue microstructure and water exchange in biophysical modelling of diffusion in white matter, MAGMA 26 (4) (2013) 345–370. doi:10.1007/s10334-013-0371-x.

[87] L. Ning, M. Nilsson, S. Lasič, C.-F. Westin, Y. Rathi, Cumulant expansions for measuring water exchange using diffusion mri, The Journal of Chemical Physics 148 (7) (2018). doi:10.1063/1.5014044.

[88] R. Song, Y.-Q. Song, M. Vembusubramanian, J. L. Paulsen, The robust identification of exchange from t2–t2 time-domain features, Journal of Magnetic Resonance 265 (2016) 164–171. doi:10.1016/j.jmr.2016.02.001.

[89] A. Ordinola, E. Özarslan, R. Bai, M. Herberthson, Limitations and generalizations of the first order kinetics reaction expression for modeling diffusion-driven exchange: Implications on nmr exchange measurements, The Journal of Chemical Physics 160 (8) (2024). doi:10.1063/5.0188865. URL 10.1063/5.0188865

[90] T. X. Cai, N. H. Williamson, R. Ravin, P. J. Basser, Multiexponential analysis of diffusion exchange times reveals a distinct exchange process associated with metabolic activity, in: Proceedings of the Annual Meeting of the International Society of Magnetic Resonance in Medicine, Vol. 31, 2023, p. 5017.

[91] E. Syková, C. Nicholson, Diffusion in brain extracellular space, Physiological Reviews 88 (4) (2008) 1277–1340. doi:10.1152/physrev.00027.2007.

[92] S. Khirevich, A. Höltzel, A. Daneyko, A. Seidel-Morgenstern, U. Tallarek, Structure–transport correlation for the diffusive tortuosity of bulk, monodisperse, random sphere packings, Journal of Chromatography A 1218 (37) (2011) 6489–6497. doi:10.1016/j.chroma.2011.07.066.

[93] M. J. Borgnia, D. Kozono, G. Calamita, P. C. Maloney, P. Agre, Functional reconstitution and characterization of aqpz, the e. coli water channel protein, Journal of Molecular Biology 291 (5) (1999) 1169–1179. doi:10.1006/jmbi.1999.3032.

[94] M. Kumar, M. Grzelakowski, J. Zilles, M. Clark, W. Meier, Highly permeable polymeric membranes based on the incorporation of the functional water channel protein aquaporin z, Proceedings of the National Academy of Sciences 104 (52) (2007) 20719–20724. doi:10.1073/pnas.0708762104.

[95] E. Solenov, H. Watanabe, G. T. Manley, A. S. Verkman, Sevenfold-reduced osmotic water permeability in primary astrocyte cultures from aqp-4-deficient mice, measured by a fluorescence quenching method, American Journal of Physiology-Cell Physiology 286 (2) (2004) C426–C432. doi:10.1152/ajpcell.00298.2003.

[96] R. Cheng, F. Zhang, M. Li, X. Wo, Y.-W. Su, W. Wang, Influence of fixation and permeabilization on the mass density of single cells: A surface plasmon resonance imaging study, Frontiers in Chemistry 7 (2019). doi:10.3389/fchem.2019.00588.

[97] D. C. Douglass, D. W. McCall, Diffusion in paraffin hydrocarbons, The Journal of Physical Chemistry 62 (9) (1958) 1102–1107. doi:10.1021/j150567a020.

[98] J. StepiŞnik, Validity limits of gaussian approximation in cumulant expansion for diffusion attenuation of spin echo, Physica B Condensed Matter 270 (1) (1999) 110–117. doi:10.1016/S0921-4526(99)00160-X.

[99] J. D. D’Errico, Slm - shape language modeling (2017). URL https://www.mathworks.com/matlabcentral/fileexchange/24443-slm-shape-language-modeling

[100] G. M. G. Shepherd, M. Raastad, P. Andersen, General and variable features of varicosity spacing along unmyelinated axons in the hippocampus and cerebellum, Proceedings of the National Academy of Sciences 99 (9) (2002) 6340–6345. doi:10.1073/pnas.052151299.

[101] S. Capuani, M. Palombo, Mini review on anomalous diffusion by mri: Potential advantages, pitfalls, limitations, nomenclature, and correct interpretation of literature, Frontiers in Physics 7 (2020). doi:10.3389/fphy.2019.00248.

[102] G. E. Uhlenbeck, L. S. Ornstein, On the theory of the brownian motion, Physical Review 36 (1930) 823–841. doi:10.1103/PhysRev.36.823.

[103] K. E. Washburn, P. T. Callaghan, Tracking pore to pore exchange using relaxation exchange spectroscopy, Physical Review Letters 97 (2006) 175502. doi:10.1103/PhysRevLett.97.175502.

[104] R. M. Henkelman, G. J. Stanisz, S. J. Graham, Magnetization transfer in mri: a review, NMR in Biomedicine 14 (2) (2001) 57–64. doi:10.1002/nbm.683.

[105] Y. Gao, B. Blümich, Analysis of three-site t2-t2 exchange nmr, Journal of Magnetic Resonance 315 (2020) 106740. doi:10.1016/j.jmr.2020.106740.

[106] B. Blümich, M. Parziale, M. Augustine, Asymmetry in three-site relaxation exchange nmr, Magnetic Resonance 4 (2) (2023) 217–229. doi:10.5194/mr-4-217-2023.

[107] Y. Scher, S. Reuveni, Y. Cohen, Constant gradient fexsy: A time-efficient method for measuring exchange, Journal of Magnetic Resonance 311 (2020) 106667. doi:10.1016/j.jmr.2019.106667.

[108] W. Warner, M. Palombo, R. Cruz, R. Callaghan, N. Shemesh, D. K. Jones, F. Dell’Acqua, A. Ianus, I. Drobnjak, Temporal diffusion ratio (tdr) for imaging restricted diffusion: Optimisation and pre-clinical demonstration, NeuroImage 269 (2023) 119930. doi:10.1016/j.neuroimage.2023.119930.

[109] E. Fieremans, L. M. Burcaw, H.-H. Lee, G. Lemberskiy, J. Veraart, D. S. Novikov, In vivo observation and biophysical interpretation of time-dependent diffusion in human white matter, NeuroImage 129 (2016) 414–427. doi:10.1016/j.neuroimage.2016.01.018.

